# Single-synapse analyses of Alzheimer’s disease implicate pathologic tau, DJ1, CD47, and ApoE

**DOI:** 10.1101/2021.06.14.448240

**Authors:** Thanaphong Phongpreecha, Chandresh R. Gajera, Candace C. Liu, Kausalia Vijayaragavan, Alan L. Chang, Martin Becker, Ramin Fallahzadeh, Rosemary Fernandez, Nadia Postupna, Emily Sherfield, Dmitry Tebaykin, Caitlin Latimer, Carol A. Shively, Thomas C. Register, Suzanne Craft, Kathleen S. Montine, Edward J. Fox, Kathleen L. Poston, C. Dirk Keene, Michael Angelo, Sean C. Bendall, Nima Aghaeepour, Thomas J. Montine

## Abstract

Synaptic molecular characterization is limited for Alzheimer’s disease (AD). We used mass cytometry to quantify 38 probes in approximately 17 million single synaptic events from human brains without pathologic change or with pure AD or Lewy body disease (LBD), non-human primates (NHP), and PS/APP mice. Synaptic molecular integrity in humans and NHP was similar. Although not detected in human synapses, Aβ was in PS/APP mice synaptic events. Clustering and pattern identification of human synapses showed expected disease-specific differences, like increased hippocampal pathologic tau in AD and reduced caudate dopamine transporter in LBD, and revealed novel findings including increased hippocampal CD47 and lowered DJ1 in AD and higher ApoE in AD dementia. Our results were independently supported by multiplex ion beam imaging of intact tissue.

## INTRODUCTION

The anatomical basis of cognitive impairment in Alzheimer’s disease (AD) is loss of limbic and neocortical synapses. This is supported by ultrastructural and microscopic studies that assessed hundreds of human synapses but with limited molecular characterization, and by molecular characterization of bulk preparations of synaptic particles from specialized brain homogenates called synaptosomes (*1–9*). More recently, array tomography (3 to 5 antibody probes) and conventional flow cytometry (2 to 4 antibody probes) have characterized hundreds of thousands to a million individual human synaptic events from synaptosome preparations (*10–12*). SynTOF (Synaptometry by Time of Flight) (*13, 14*) quantifies tens of millions of individual synaptic events by mass cytometry, permitting robust machine learning (ML) applications. Here, we performed SynTOF on healthy non-human primates, PS/APP mice, and human synaptosomes from carefully selected individuals with only AD neuropathologic change (ADNC) or only Lewy body disease (LBD) (*15, 16*), as well as normal age-matched adults. We tested the hypothesis that the synaptic molecular composition in humans with only AD resembled aged PS/APP mice, one of the most widely used transgenic models of AD. We then used ML approaches to identify presynaptic subpopulations in controls, and to discover novel disease-related changes in synaptic molecular composition. Finally, we confirmed the major features of ML-identified synaptic subpopulations with multiplexed ion beam imaging (MIBI).

## RESULTS

### Generation of single synaptic data via SynTOF

Over five years, 113 synaptosome preparations were made from postmortem human brain collected from carefully annotated adult research participants, 21 of whom met rigorous inclusion/exclusion criteria for both clinical and pathologic features: Controls (n=6), high level AD neuropathologic change (ADNC) only (n=9), and high level LBD only (n=6) (**Table S1**); note that the ADNC and LBD groups each contained two resilient individuals, meaning high level pathologic change without clinical diagnosis of dementia and/or Parkinson’s disease (*17*). Following our established protocol (*14*), samples were prepared, barcoded, stained with a 38-plex SynTOF panel (**Table S2**), and analyzed to detect approximately 100 million events from human synaptosome preparations. Debarcoding and sequential gating (*14*) yielded highly enriched single human pre- (n=14,904,100) and postsynaptic (n=823,280) events from Brodmann area (BA) 9 of prefrontal cortex, caudate nucleus, and hippocampus (synaptosome preparations strongly favor pre- over postsynaptic particles). Using the same method and multiplex panel, we also detected >740,000 single presynaptic events from non-human primate (NHP) frontal cortex and striatum, and >440,000 single presynaptic events from cerebral cortex and hippocampus of 22 month-old wild type C57Bl6 and PS/APP transgenic mice (*18*). These 38-dimension intensity data from approximately 17 million single synaptic events comprise the data analyzed.

### Pseudo-bulk analysis of human and mouse synaptic events

The first goal was to analyze the data by established gating protocols (*13, 14*) to facilitate comparison with existing human data obtained using other technologies, and to compare across species. Control human and NHP synaptosomes were similar with few significant differences between primate species (**Fig. S1**) (*19*), underscoring the molecular integrity of human synaptosomes as prepared and analyzed here. Expected regional variation was observed in Control presynaptic events (Fig. 1A), including reciprocal variation in average glutamatergic (P<0.01) and GABAergic (P<0.05) events between BA9 and caudate, and greater % DAT positive presynaptic events in caudate vs. BA9 (P<0.001) or vs. hippocampus (P<0.001) (**Fig. S2**) (*20–25*). Pre-/post-synaptic ratio in Controls for proteins known to be enriched in the presynapse ranged from 5-fold to >200-fold (Fig. 1B). By SynTOF, presynaptic average % positive events were 4.3±0.4% for Aβ40 and 2.0±0.2% for Aβ42 in BA9, and 18.1±1.7% for PHF-tau in hippocampus for ADNC; these results align well with 0.6% to 5.1% Aβ positive (*10*) and 15.4% pathologic tau positive (*12*) presynaptic synaptosomes in AD by array tomography. Our results demonstrate that human SynTOF data derive from synaptic preparations with high molecular integrity, show expected patterns of anatomical distribution, and compare well with data collected by others using transmission electron microscopy or array tomography.

**Fig. 1.**
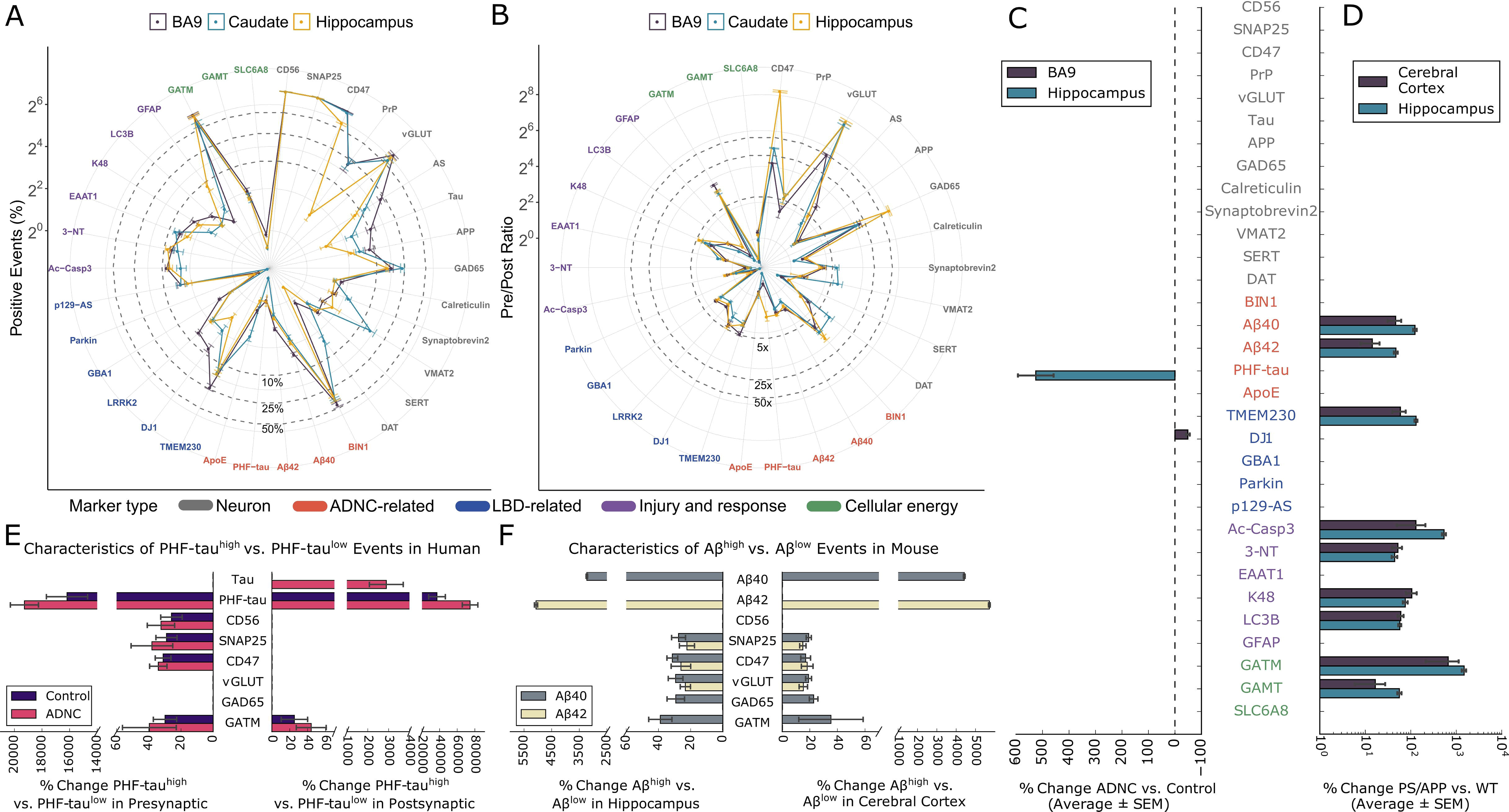
SynTOF panel results for human and mouse synaptosomes analyzed as average % positive events (±SEM). (**A**) Spider plot of % positive events from BA9, caudate nucleus, and hippocampus from human Control (n=6). (**B**) Spider plot of presynaptic/postsynaptic event ratios for non-zero postsynaptic markers. (**C**) Presynaptic average % change in % positive events (±SEM) in BA9, caudate, and hippocampus from human Control, ADNC (n=9), and LBD (n=6) analyzed by two-way ANOVA with brain region vs diagnostic group. Only results with significant differences (P<0.001) are presented. Results for LBD samples are presented in text. (**D**) Similar to (**C**) but for cerebral cortical and hippocampal presynaptic results from 22 month-old PS/APP (n=5) vs. WT (n=5) mice. (**E**) Hippocampus presynaptic and postsynaptic % change in mean intensity (±SEM) in both Control and ADNC samples analyzed by two-way ANOVA with PHF-tau^high^ or PHF-tau^low^ vs. mean intensity for each probe. Only significantly greater mean intensity following correction for multiple comparisons are presented. (**F**) Similar to (**E**) but for hippocampus or cerebral cortex presynaptic events in PS/APP mice divided into Aβ40^high^ vs. Aβ40^low^ events or Aβ42^high^ vs. Aβ42^low^ events.

Presynaptic average % positive events for two probes were significantly different (P<0.001) when comparing ADNC to Controls: hippocampus had more PHF-tau (525%) and BA9 had less DJ1 (49%) (Fig. 1C). LBD was significantly different from Control only for 51% reduction in caudate DAT^high^ presynaptic events (P<0.001), aligning closely with estimates from biochemical studies and SPECT imaging (*26*). Importantly, lack of significant presynaptic changes in LBD BA9 and hippocampus underscored that the changes in AD were disease-specific and not merely covariates of neurodegeneration or dementia such as debilitation or reduced mobility. Although previous studies identified an approximately 20% increase in p129-α-synuclein positive presynaptic synaptosomes in cingulate gyrus and putamen from LBD samples (*11*); we did not observe this in the three regions analyzed here. Postsynaptic average % positive PHF-tau also was significantly increased in ADNC hippocampus compared to Controls (P<0.0001, % change = 320±51), again in agreement with array tomography (*12*).

We tested the hypothesis that Aβ-bearing presynaptic events could be detected by SynTOF by using a common transgenic mouse model of AD. Cerebral cortical and hippocampal synaptosomes from 22 month-old WT (n=5) and PS/APP (n=5) mice were analyzed by SynTOF exactly as above. Eleven probes had significantly increased presynaptic average % positive events in PS/APP compared to WT mice (P<0.001, Fig. 1D); no probe’s signal was significantly decreased. By SynTOF, aged PS/APP mice had approximately 100-fold increase in presynaptic Aβ accumulation as well as many of the responses to Aβ-induced injury that have been reported previously in bulk tissue from AD transgenic mouse models, including: increased ubiquitin conjugation (K48), lysosomal stress (LC3B), increased free radical injury (3NT), activated apoptotic signaling (Ac Casp3), and cellular energy stress as indicated by significantly increased presynaptic expression of the two enzymes that catalyze creatine synthesis (GATM and GAMT, Fig. 1D) (*27, 28*). An unexpected finding was increased transgenic mouse presynaptic TMEM230, a protein product of a Parkinson’s disease risk gene, in aged PS/APP mice. Human hippocampus presynaptic PHF-tau^high^ events, whether from ADNC (both dementia and resilient) or Control, yielded similar results indicating a stress response to neurofibrillary change independent of clinical status. This included increased capacity to synthesize creatinine in both pre- and postsynaptic events, as well as increased expression of multiple presynaptic proteins, including CD56 and CD47, perhaps as attempts to forestall degeneration (Fig. 1E). Human hippocampus PHF-tau^high^ postsynaptic events again had increased expression of GAMT in Controls and ADNC, and increased tau expression only in ADNC, perhaps indicating an attempt to mitigate sequestration of pathologic tau. Hippocampus and cerebral cortex presynaptic events from aged PS/APP mice were divided into Aβ42^high^ vs. Aβ42^low^ or Aβ40^high^ vs. Aβ40^low^ events and mean intensity of each probe analyzed; there were too few mouse postsynaptic events to analyze confidently (Fig. 1F). Interestingly, Aβ42^high^ and Aβ40^high^ events were largely distinct populations. In both regions, Aβ42^high^ or Aβ40^high^ events shared significantly increased (P<0.001) expression of vGLUT, SNAP25, and GATM (except cortex for Aβ42^high^). Aβ40^high^ presynaptic events tended to have higher GAD65 expression in both regions (P<0.01). Therefore, pseudo-bulk analyses validated SynTOF data by close agreement with results obtained by other methods in humans and transgenic mice.

### Identification of subpopulations from human presynaptic events

We adopted a clustering algorithm based on modified autoencoder (AE) neural networks to identify subpopulations from single synaptic events. Briefly, the AE was used to learn compressed hidden representations of the single synapse data, then a K-Mean-based soft clustering layer was appended and trained with AE to cluster the compressed representation. Using a subset of 24 designated phenotypic markers that did not include disease-specific markers (**Table S2**), the modified AE clustered Control single presynaptic events from all three brain regions into 15 different subpopulations (see details in **Methods**). Figure 2A shows each subpopulation’s standardized mean expression for each probe in all brain regions for Control samples; these characteristics remained similar when stratified by brain region (**Fig. S3A**). Subpopulation A comprised two “high-expresser” subpopulations (A1 and A2) with high signals from most probes. In contrast, subpopulation B contained two subpopulations (B1 and B2) with low signal from most probes, with the exception of GATM and GAMT. Subpopulation C’s eleven subpopulations were heterogeneous; C1 to C5 tended to have moderately higher probe signals than C6 to C11. Several C subpopulations had unique features, *e.g*., C3 and C6 were GAD65^high^, C4 was PrP^high^, C8 was VGLUT^high^, and C10 was GAMT^high^. Single presynaptic event data visualized by t-SNE dimensionality reduction showed proximity among subpopulations (Fig. 2B), and their characteristics were appreciated by individual phenotypic marker expressions (**Fig. S4**). Figure 2B also visualizes select subpopulations, such as BA9’s A1, B1, and C4, by density plots of a 2-step gating strategy optimized by GateFinder (*29*).

**Fig. 2.**
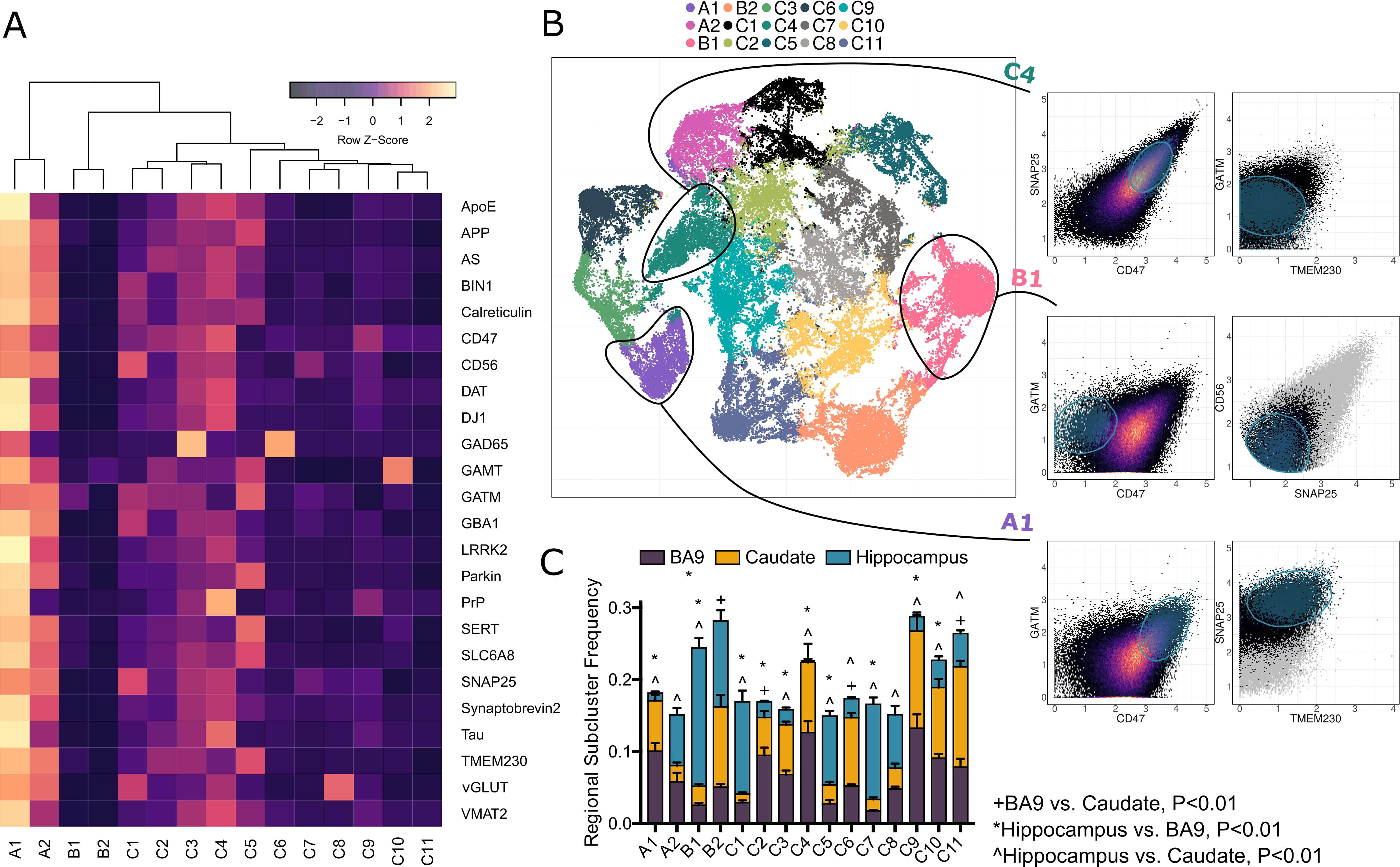
Modified deep autoencoder clustered the presynaptic events from Control samples to different subpopulations according to the designated phenotypic markers. (**A**) A heatmap showing the mean expression values from Control samples (all brain regions) for each of the phenotyping markers across all identified subpopulations. The mean expression values were scaled across subpopulations for differential visualization purposes. (**B**) A t-SNE plot of the autoencoder’s hidden representations of presynaptic events randomly selected from Control samples across all brain regions colored by cluster association (left), and biaxial plots of gating strategy obtained from GateFinder to achieve the selected examples of subpopulations from a sample of BA9 region (A1, B1, and C4). The black-to-yellow color scale represents the density of the events, while the gray represents events that were excluded by the previous gating step. (**C**) The subpopulation’s mean frequency (±SEM) from Control samples in BA9, caudate nucleus, and hippocampus with symbols indicating Wilcoxon’s P<0.01 after Tukey’s correction.

Subpopulation frequency varied significantly across brain regions (P<0.0001) (Fig. 2C; **Fig. S5A** to **C**). BA9 was similar to caudate in most subpopulations, while hippocampus tended to be different from the other two regions. Corrected multiple comparisons revealed significantly (P<0.01) different patterns of regional subpopulation frequencies: BA9 was different from caudate in B2, C2, C6, and C11; BA9 was different from hippocampus in A1, B1, B2, C1 to C5, C7, C9, and C10; and caudate was different from hippocampus in A1, A2, B1, C1, and C3 to C11. These results show a relatively distinct frequency of Control presynaptic subpopulations in the human hippocampus. Since B-type subpopulations were the most frequent in hippocampus, our results suggest that subpopulation Bs were mostly undifferentiated subpopulations based on our selected markers, and that further refinement of our SynTOF panel may be needed to reflect more fully hippocampus presynaptic diversity.

### Presynaptic subpopulations confirmed pseudo-bulk findings and revealed additional strong signals from ADNC and LBD presynaptic events

Based on the subpopulations identified in Fig. 2A, we next focused on using conventional ML to identify regional differences in presynaptic molecular composition among the three groups: Control, ADNC, and LBD. Specifically, mean expression for the full complement of presynaptic markers was calculated for each subpopulation, resulting in 1530 expression features per participant (3 brain regions × 15 subpopulations × 34 probes; note that 4 probes were excluded because they were used for gating). Distinct from the earlier pseudo-bulk analyses, features with complete separation between the two groups, which would result in perfect classification (**Table S3**), were intentionally omitted in ML models to allow identification of other less strong features. From the six ML algorithms we chose for comparison (see details in **Methods**), an elastic net (EN) model outperformed others, achieving an AUC of 0.96 with significantly different EN predicted values for Control vs. ADNC (P=1.6×10^-3^) (Fig. 3A); this was despite the exclusion of features with complete separation between groups, which were hippocampus PHF-tau from most subpopulations and hippocampus CD47 from six subpopulations (**Fig. S6**). Notably, the Control sample with the highest EN predicted probability value (0.37) was the only Control individual who had been diagnosed previously as MCI, but at final consensus was classified as cognitively normal. FDR-adjusted P values (Q values) of each feature alongside EN’s weights (Fig. 3B) confirmed pseudo-bulk analysis by showing increased hippocampus PHF-tau mean intensity and reduced BA9 DJ1 mean intensity significantly separated ADNC from Control in almost every subpopulation (Fig. 3C and D, **Fig. S7**). An additional strong feature was increased hippocampus CD47 mean expression in ADNC, particularly in CD47^high^ subpopulations (Fig. 3E, **Fig. S7**), which also tended to express higher PHF-tau (Spearman’s R=0.62, P<<2.2×10^-16^). Other markers exhibiting strong univariate associations with ADNC (Q values <0.05) or associated with high EN coefficients, but less prevalent across subpopulations, such as GATM and K48 in caudate and 3-NT in BA9, are summarized in **Fig. S7**.

**Fig. 3.**
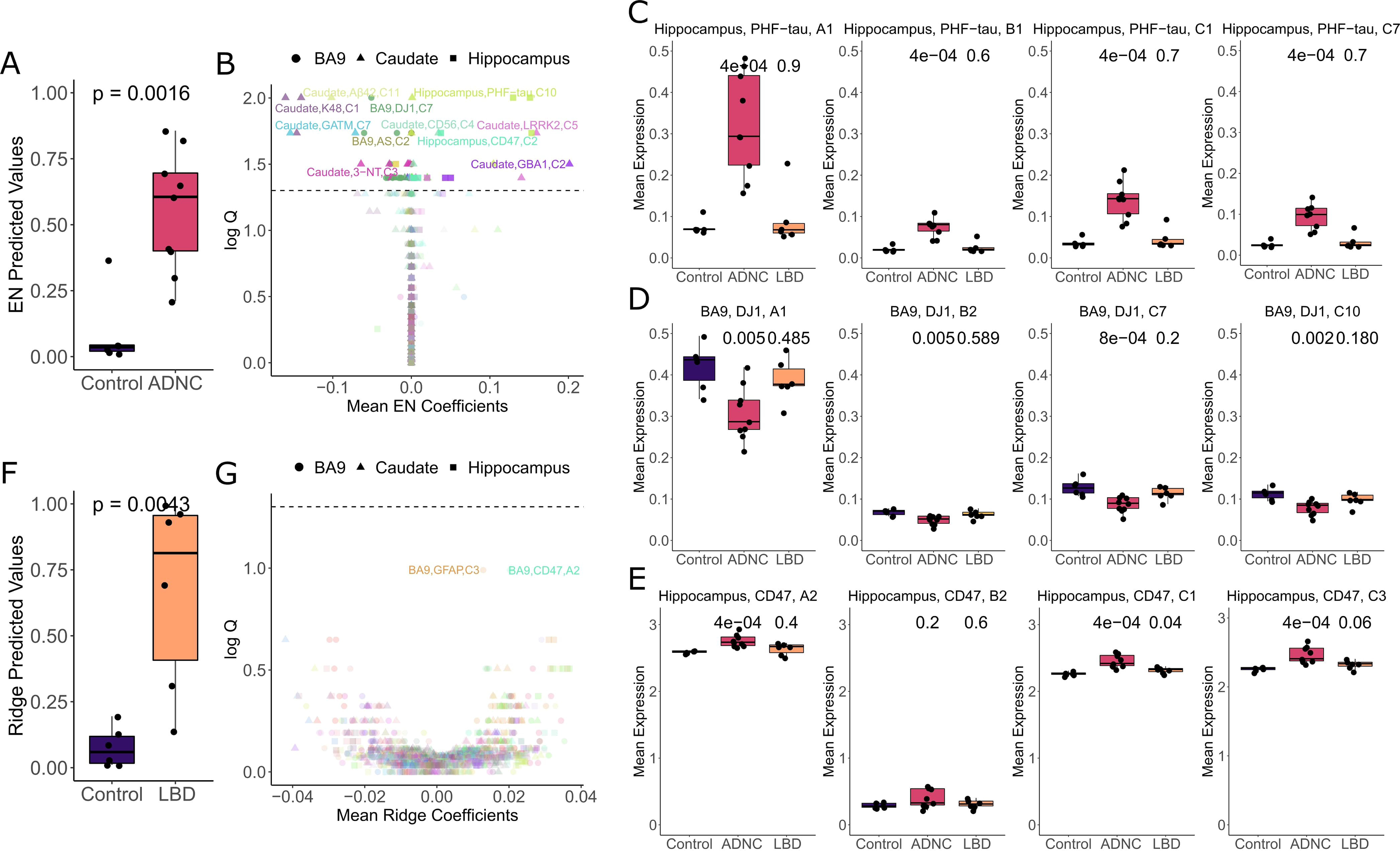
Machine learning model and univariate analyses of the subpopulations identified more predictive signals for Controls vs. ADNC and LBD. (**A**) The predicted values from elastic net model on Control vs. ADNC. In the model, features with complete Control vs. ADNC separation, such as most PHF-tau and some CD47 signals in hippocampus subpopulations, were intentionally removed to allow observation of predictive performance of other features (**Table S3**). (**B**) Visualization of important features through EN model’s coefficients coupled with univariate Q values (from FDR-corrected Spearman’s P values). The dashed line represents a Q value of 0.05. The color corresponds to each of the expression marker types and the shape indicates brain region. (**C**)-(**E**) Boxplots of selected markers whose signal intensity was significantly different across multiple subpopulations for ADNC, including PHF-tau, CD47, and DJ1 signals from one A and B subpopulations and two C subpopulations. Note that in this panel only CD47 expression in the hippocampal subpopulation B2 was not significantly different, but was selected as a counter-example to support that CD47 differences were found in CD47^high^ subpopulations only. (**F**)-(**G**) Similar to (**A**)-(**B**) for Control vs. LBD samples.

A ridge regression model outperformed other ML approaches for Control vs. LBD, achieving an AUC of 0.97 and ridge predicted values (P = 4.3×10^-3^) (Fig. 3F), even with exclusion of features with complete separation, which were lower caudate DAT from most subpopulations and higher BA9 GFAP from subpopulations A1 and C4 in the LBD group (**Table S3**). In both of these subpopulations significantly increased signals from the other astrocytic marker, EAAT1, also were observed albeit weaker (data not shown). Severe astrogliosis, even with spongiform change, in cerebral cortical superficial layers is a well described feature of LBD but not AD (*15, 30*). Similar to ADNC, features associated with greater ridge coefficients were those with higher univariate impact (Fig. 3G); unlike ADNC, none of the features passed the 0.05 Q value threshold, except for the excluded features that completely separated the two groups (**Table S3**). This may be due in part to a lower number of LBD only samples, or perhaps less severe degeneration in LBD compared to ADNC (*31*). Notably, none of the top features for Control vs. LBD, *i.e*., caudate DAT in multiple subpopulations, BA9 GFAP in C3, and BA9 CD47 in A2, overlapped with those from Control vs. AD, again indicating that SynTOF identified disease-specific presynaptic molecular features.

### Presynaptic features suggest molecular differences between AD dementia and AD resilient cases

Resilience describes maintained cognitive function despite high levels of pathologic change. There were two AD resilient cases among the nine ADNC analyzed, and also the only ones out of the entire original set of 113 samples (**Table S1**). To examine their impact, these two AD resilient cases were sequestered and Q values recalculated. The analysis (6 Control vs. 7 AD dementia) resulted in robust increase in existing signal strength and prevalence, particularly for DJ1 from more subpopulations in BA9 (Fig. 4A and **Fig. S7**), except for PHF-tau and CD47 which remained the same. Several new signals also emerged with the strongest being ApoE from subpopulations A1 and C5 in hippocampus (Fig. 4B). Further gating to obtain GFAP-EAAT1-presynaptic events, *i.e.* presynaptic events without attached astrocytic remnants, removed 11.4±3.1% of presynaptic events. Analyses of the GFAP-EAAT1-presynaptic event data suggested that the ApoE signal derived from neuronal elements with and without attached astrocytic remnants (Fig. 4C). No new signal was revealed when a similar approach was used with LBD resistant cases (not shown).

**Fig. 4.**
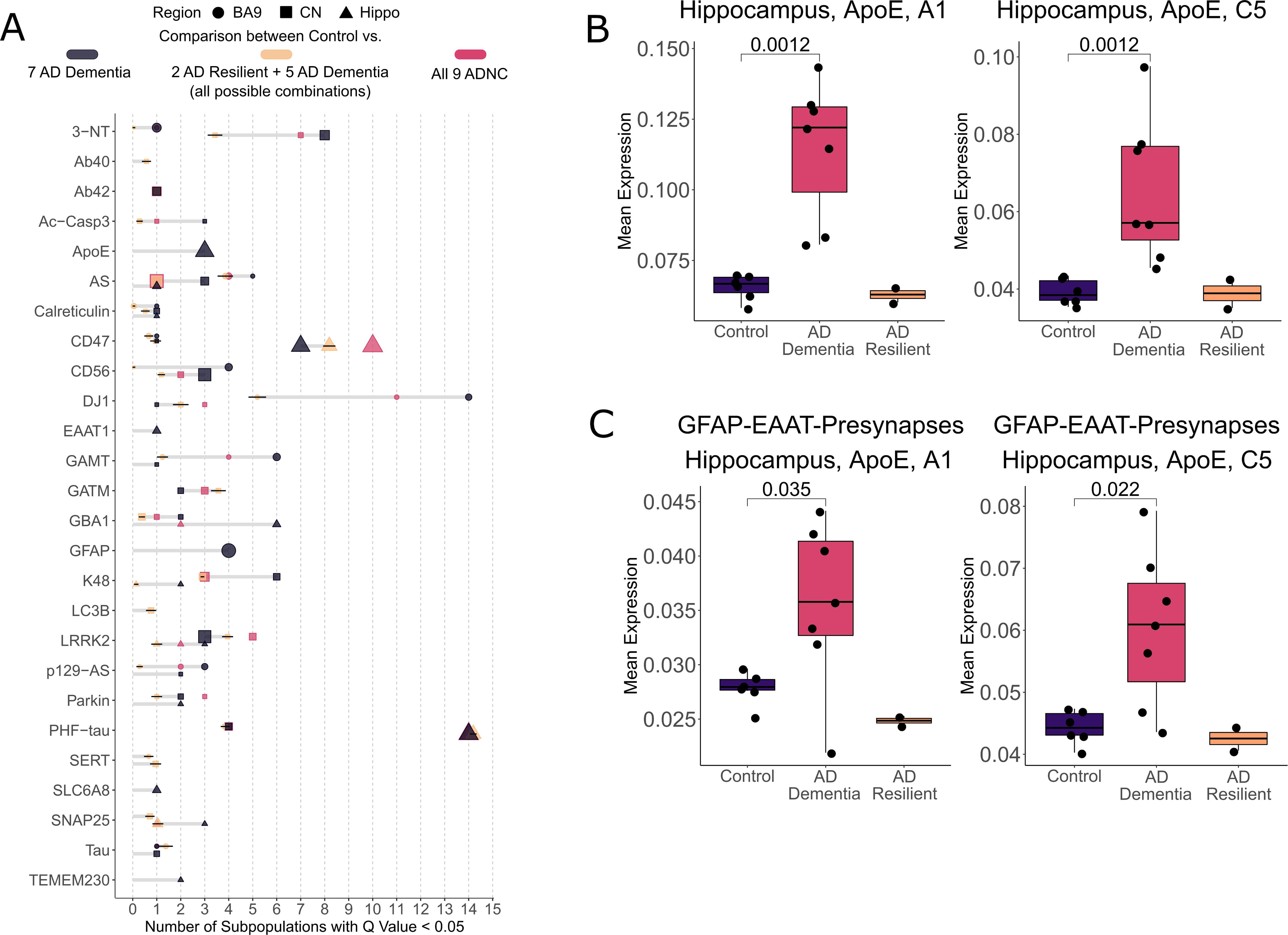
Changes in significant features and signal strength within AD dementia cases. (**A**) The number of significant subpopulations after FDR adjustment (Q value <0.05) for each of the expression markers in different brain regions. The color denotes comparison type. Dark gray indicates the comparison between 7 AD dementia cases and all Control cases. Tan is between every possible combination (n=36) of all Control cases and 7 ADNC cases (all possible combinations of 5 out of 7 AD dementia plus the 2 AD resilient) therefore, the purple dots represent the mean and standard deviation for these 36 combinations. For reference, the red color indicates the original comparison between all 9 ADNC cases and the Control. The size of the dots is proportional to the mean Q value of all significant subpopulations, where larger size indicates lower Q value. Only markers with significant subpopulations are shown. (**B**) Boxplots of the strongest, newly emerged signals for Control vs. AD dementia only cases:ApoE expression from 2 subpopulations in the hippocampus. The plot separately visualizes Control, AD dementia and AD resilient cases. **c**, Data in subfigure (**A**) & (**B**) include the approximately 11% of events that contained attached astrocytic remnants. To assess the extent to which ApoE expression derived from neuronal events only vs. the subset of presynaptic events with astrocytic remnants, we repeated our analysis following gating (GFAP- and EAAT1-) that removed the small subset of events with astrocytic signal. GFAP-EAAT1-presynaptic events had reduced effect size compared to all presynaptic events (subfigure (**B**)), but still remained significant, suggesting that the ApoE signal derived from both neuronal presynapse and astrocytic remnant components of the single event data.

To support that these new signals obtained upon removal of two AD resilient cases were likely not coincidental, an additional analysis was conducted wherein the two AD resilient cases were retained and instead two AD dementia cases were removed and Q values calculated. No significant change in marker signals, including ApoE, occurred with sequestration of any other pair of AD dementia cases (Fig. 4A). Although limited by a small number of rare cases, our analyses suggested that the amount of hippocampal ApoE in the presynapse, tentatively in both the neuronal component and intimately associated astrocytic component, may be important to the clinical expression of dementia in AD.

### MIBI-TOF analysis of tissue sections supports SynTOF findings

Multiplexed Ion Beam Imaging by Time-of-Flight (MIBI-TOF) (*32*) was used to support the physical existence of autoencoder-identified subpopulations. Akin to SynTOF, elemental isotopic reporters of n-dimension are used for quantitative profiling, but MIBI-TOF also determines nanometer spatial resolution. A 2.4 mm^2^ region of interest (ROI) composed of six 400 μm^2^ fields of view (FOVs; 1024×1024 pixels each) from the CA1 sector of Control and ADNC hippocampus were scanned at 300 nm resolution (Fig. 5A). Figure 5B shows example FOVs at different scales colored for synaptophysin, CD47, and PHF-tau where the zoomed-in images suggest close proximity of some PHF-tau and CD47 pixels. Validation of SynTOF data, derived from physically enriched pre-synaptic particles, by MIBI-TOF requires *in silico* enrichment of presynapse signals vs. all other signals in the tissue section; this was achieved by co-locating at least two presynaptic markers. First, each pixel was agnostically assigned as positive or negative for each marker through self-organizing map clustering (Fig. 5B, see details in **Methods**); MIBI-TOF marker positivity is equivalent to high or very-high expression in SynTOF. Second, a sliding window collected data over the entire MIBI-TOF FOV for quantitative analysis, where windows that contained CD47+Synaptophysin+PSD95-pixels were defined as presynaptic. A 6×6 pixel (approximately 1.8 μm^2^) window was used because a presynapse is approximately 1 μm^3^ in fixed and embedded tissue (*33*).

**Fig. 5.**
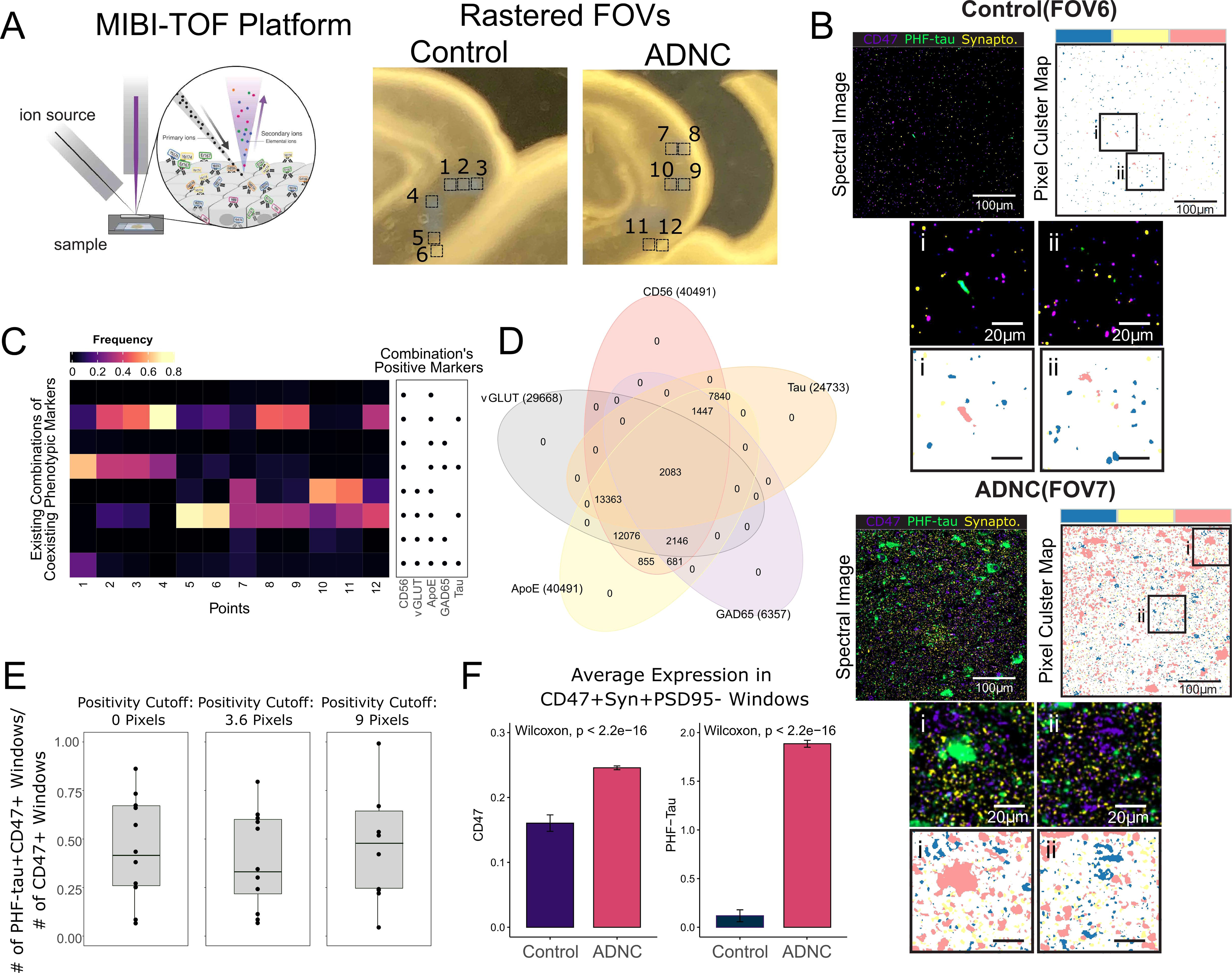
Analysis of MIBI-TOF images supports SynTOF findings by suggesting high coexistence of phenotypic markers, coexistence of CD47 and PHF-tau, and higher CD47 expression in ADNC. (**A**) Illustration of MIBI-TOF and the sampled locations of the FOVs from CA1 sector of the hippocampus. (**B**) Examples of MIBI-TOF image FOV from Control and ADNC. The spectral images are overlaid with CD47, PHF-tau, and synaptophysin channels for identification of likely synapses (left) with corresponding areas colored by CD47 (blue), PHF-tau (pink), and synaptophysin (yellow) positive clusters obtained from SOM clustering (right). Bottom, smaller panels show zoomed-in images within the rectangles. (**C**) A heatmap showing existing combinations of positive MIBI’s phenotypic markers found from aggregated sliding windows with colors representing a fraction of positivity over the number of presynaptic windows defined as containing CD47+Synaptophysin+PSD95-. (**D**) A Venn diagram summarizing the intersected expression of the phenotypic markers. The number in the parenthesis indicates the total number of positive windows. (**E**) Boxplots showing the fraction of PHF-tau and CD47 coexistence within a window over different numbers of pixel cut-offs used for defining marker positivity. (**F**) Barplots showing higher average (±SEM) CD47 and PHF-tau expressions in ADNC based on all presynaptic windows in all points.

MIBI-TOF confirmed three major findings from SynTOF. First, MIBI-TOF displayed high coexistence of multiple positive markers in each sliding window, in accord with subpopulation characteristics identified by SynTOF, particularly the “high-expressers” such as subpopulation A. Indeed, a heatmap and Venn diagram (Fig. 5C & D) of a subset of SynTOF phenotypic markers used in MIBI-TOF (ApoE, CD56, GAD65, tau, and vGLUT) showed high frequency of coincident marker positivity in presynapse windows. Second, presynaptic colocation of CD47 and PHF-tau observed in SynTOF also was observed by MIBI. Figure 5E shows that about half of the CD47+ presynaptic windows also had PHF-tau+ pixels. This result was consistent even if the positivity cut-off for the number of pixels was increased. Finally, the average level of expression of PHF-tau and CD47 were both higher in ADNC presynaptic windows than in Control, again validating SynTOF results (Fig. 5F). All analyses were also repeated using 3×3 pixel and 9×9 pixel window sizes and arrived at the same conclusions (**Fig. S8**). In aggregate, using a complementary method on intact tissue, MIBI-TOF data confirmed the major findings from SynTOF.

## DISCUSSION

The anatomical basis for cognitive decline in AD and LBD is synaptic injury. SynTOF has advantages over other techniques that molecularly characterize synaptic events from synaptosome preparations: (i) further computational enrichment of single synaptic events by error correction and sequential gating, (ii) greater than an order of magnitude increases in detection of enriched synaptic events, and (iii) an order of magnitude increase in number of multiplex probes. We built our SynTOF panel to focus on phenotypic markers, products of risk genes, and mechanisms relevant to AD or LBD. As shown previously with conventional flow cytometry (*19*) and replicated here with SynTOF, human synaptosomes maintain molecular integrity at least as compared to NHP samples collected and prepared under optimal research conditions. Results from our SynTOF panel aligned well with ultrastructural estimates of the regional proportions of synaptic subtypes and with array tomography estimates of synaptic involvement by hallmark pathologic proteins. Furthermore, MIBI-TOF colocalization of prominent molecular features supported the major outcomes of autoencoder-identified subpopulations. Thus buttressed, SynTOF unveiled key differences in presynaptic molecular composition between late-onset AD and aged PS/APP mice, reinforced the relevance of synaptic pathologic tau species in AD (*34*), and highlighted presynaptic roles for CD47, DJ1, and ApoE in AD.

Mutations in *PARK7*, the gene that encodes DJ1, cause an autosomal recessive form of early onset PD, and for this reason was included in our SynTOF panel. DJ1 functions as a redox sensitive chaperone and antioxidant. It is a target for reversible and irreversible oxidative modification in brains of patients not only with PD but also with late-onset AD where it may incorporate in hallmark lesions like Lewy bodies and amyloid plaques (*35, 36*). These pathologic changes are proposed to underlie the significant accumulation of DJ1 in the detergent-insoluble fraction of bulk tissue from affected regions of the brain in AD and LBD (*35, 37, 38*). Our results showed significantly reduced BA9 presynaptic DJ1, which is consistent with these findings from others, and supports the hypothesis that increased oxidative stress can lead to post-translational modifications of synaptic DJ1 that result in its shift to non-synaptic compartments. We observed significant reduction in presynaptic DJ1 only in AD, consistent with earlier data showing that oxidative damage is greater in AD than in LBD and other related dementias (*39*).

CD47 is localized to dendrites of cultured neurons and acts as the presynaptic receptor for the shed ectodomain of signal regulatory protein-alpha (SIRP-alpha) in activity-dependent trans-synaptic maturation of functional synapses (*40, 41*). CD47 also is expressed on myeloid cells, and has been shown to impact microglial and macrophage activity in mouse models of multiple sclerosis and cell culture models using Aβ peptides (*42, 43*). Importantly, CD47, an immune ‘don’t eat me signal’, protects from excess microglia-mediated synaptic pruning during mouse development (*44*), but as far as we are aware there has been no investigation of synaptic CD47 in brain aging or neurodegenerative disease. Our results showed that presynaptic CD47 was significantly elevated in AD hippocampus, with less robust increases in LBD hippocampus or BA9 for both diagnostic groups; caudate presynaptic CD47 was not different among the three primary diagnostic groups. Since activated microglia-mediated neuronal injury is now a well established mechanism in AD (*45*), we hypothesize that increased presynaptic CD47 is a response to limit or counteract synaptic degradation by microglia in hippocampus and likely cerebral cortex. Our discovery of elevated CD47 in AD hippocampus presynaptic events, which also tended to coexpress with PHF-tau, is especially encouraging because of emerging CD47-targeting therapeutics in certain forms of cancer (*46*).

There is intense interest in where resilience may reside in the cascade of events from synaptic injury to symptoms because this may illuminate effective therapeutic targets. Our results suggest that resilience to AD may be related to lower ApoE levels in presynaptic events, either in neuronal components or attached associated astrocytic remnants. Multiple experimental models support this proposal at the tissue level. For example, *ApoE* deletion or haploinsufficiency reduces cerebral Aβ amyloidosis and rescues from tau-induced neurodegeneration (*47, 48*). Moreover, approximately 50% reduction of brain ApoE in a humanized transgenic mouse model of AD reduced neuronal injury perhaps from suppressed glial response from lowered ApoE levels (*49*). Although limited by a small number of rare resilient cases, our results from human presynaptic events align with multiple experimental models to suggest that lower levels of presynaptic ApoE are a significant component of the resilient state.

Our pseudo-bulk analysis showed that only a small subset of human presynaptic events had detectable Aβ peptides, similar to previous work (*10*), and that synaptic Aβ peptides were not increased in AD. We showed that synaptic particles with accumulated Aβ peptides survive synaptosome preparation and downstream analysis by investigating aged PS/APP mice for which there is a large amount of experimental data supporting increased synaptic Aβ peptides (*27*). Importantly, the pattern of presynaptic injury and response in PS/APP mice was different from human AD samples. It is possible that the transgenic mice reflect a different stage of disease than humans with sporadic AD (*28*), but this is unlikely the sole explanation for the differences in presynaptic changes because the brain autopsies were from people with high level ADNC at all stages of cognition, a rough proxy for extent of neurodegeneration. A more direct comparison to PS/APP mice would be individuals with *APP* or *PSEN1* mutations; however, as far as we are aware, no appropriately prepared tissue from individuals with these disease-causing mutations exists in the world. Together, our results support a role for Aβ peptide-mediated presynaptic injury in PS/APP mice, but caution that there is a critical difference in the synaptic distribution of Aβ peptides between aged PS/APP mice and people with sporadic, high level ADNC.

In summary, SynTOF provides an unparalleled opportunity for multiplex analysis of millions of synaptic events. Results from our current panel of 38 probes implicated pathologic tau, immune-mediated pruning, and oxidative injury, but not increased Aβ peptide accumulation, as key components of synaptic injury in AD, and synapse-associated ApoE as important in the clinical expression of dementia.

## MATERIALS AND METHODS

### Study design

The research goal is to investigate synaptic molecular changes in brain samples from individuals with ADNC and LBD. All brain donations were from research volunteers at Stanford University or the University of Washington. All participants provided informed consent approved by Institutional Review Boards. Clinical and pathological diagnoses followed consensus criteria (*15, 30, 50*). One hundred thirteen samples were collected and cryopreserved prospectively from brain autopsies with short post-mortem interval (PMI < 8 hr) at the end of 5 years; only those that met stringent clinical and pathological criteria for Control (n=6), ADNC (n=9), and LBD (n=6) were used for this study (**Table S1**) (*13, 19*). Samples were processed and mass synaptometry data were acquired, barcoded/debarcoded, and gated for % positive events exactly as described by us (*14*) with the exception of adding gephyrin to PSD95 to gate for postsynaptic events. No samples were regarded as outliers or excluded.

Animal models, PS/APP (n=5) and C57Bl6 wild type (n=5) male mice, were included to evaluate Aβ-containing synaptosomes and for comparison with humans. All mouse procedures were conducted in accordance with the guidelines of the Institutional Animal Care and Use Committee at Stanford University. Mice were sacrificed at 22 months of age, their brains rapidly removed, and brain regions collected for synaptosome preparation as previously described (*13, 19*). Additionally, brain regional synaptosomes from female Macaca fascicularis (n=4) collected under research protocols were used to assess the integrity of human synaptosomes collected through rapid autopsy. All non-human primates (NHP) procedures were performed at Wake Forest University (Winston-Salem, NC) with approval from the Institutional Animal Care and Use Committee, according to recommendations in the Guide for Care and Use of Laboratory Animals (Institute for Laboratory Animal Research), and in compliance with the US Department of Agriculture Animal Welfare Act and Animal Welfare Regulations (Animal Welfare Act as Amended; Animal Welfare Regulations). NHP were sacrificed at 11.5±0.3 years of age, brains were immediately removed, and regions collected for synaptosome preparation using the same human protocol as previously described by us (*19*). NHP samples were prepared for mass cytometry and data acquired and processed exactly as for human samples (*14*).

### Clustering of single-synaptic events

Cellular population assignment through clustering algorithms has developed rapidly in recent years (*51–54*). In our model, an autoencoder (AE) was used to cluster single-synaptic events to different subpopulations. AE is a type of neural network typically with a bottleneck layer that generates compressed representations of the single-synapse data through learning to reconstruct them. Our model is adapted from a variant of AE called Improved Deep Embedded Clustering (IDEC) (*55*), which adds to typical AE a clustering network layer that clusters single-synapse data based on the obtained compressed representation. This method was selected over traditional single-event clustering algorithms, such as FlowSOM, SPADE, or k-means, due to its higher cluster assignment accuracy compared to benchmarked manual gating on publicly available datasets.

In our model, two identical autoencoders with different numbers of bottleneck nodes were employed. The node sizes for each layer were [512, 256, 128, 10, 128, 256, 512] and [512, 256, 128, 5, 128, 256, 512]. The autoencoders were separately pre-trained to reconstruct the single presynaptic events from all brain regions of all 6 Control samples using mean square error (MSE) for loss function and adaptive gradient method for optimization (Adagrad). After pre-training, a clustering layer was appended to the autoencoders where its input was a concatenated hidden representation from the two bottleneck layers. The weights of the clustering layer were initiated using centers from K-means clustering. Its number of nodes, which dictated the number of clusters, was determined using an elbow method, *i.e.* the number of clusters at which the within-cluster-sum of squared errors (WSS) plateaus. This new architecture was then trained to minimize the following loss function:

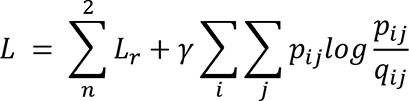

where the first term is the summation of MSE loss for reconstruction of the two autoencoders and the second term is the clustering assignment hardening loss with proportion modulated by *γ* of 1. The clustering loss was calculated by:

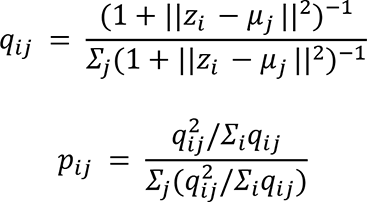

where *q_ij_* the normalized similarity between hidden representation point *z_i_* to cluster centroid *μ*_j_ by student Student’s *t*-distribution measure, and can be considered as soft cluster assignments, while *p_ij_* served as stricter (hardened) auxiliary ground truth labels on which the model trained. Therefore, *p_ij_* was not updated after every batch interval, but every 700 to allow convergent cluster layer training. The entire autoencoder model was trained until the hard cluster assignment calculated by *argmax*(*q_ij_*) became less than 5% difference compared to the last update. To leverage inherent randomness within neural networks, these steps were repeated ten times and the final cluster assignment was obtained from meta-clustering (greedy algorithm by Dimitriadou, Weingessel and Hornik [DWH]) of the hard cluster assignment obtained from each of the iterations (*56*). The trained model was then used to predict cluster assignment for post-synaptic events and presynaptic events of other diagnostic groups.

For each cluster and brain region in a given sample, a mean value for each of the 34 markers (4 of the 38 total markers were withheld because they were used in negative gating presynaptic events) was calculated, yielding a total of 1530 expression features per participant (3 brain regions x 34 markers x 15 subpopulations). Mean expression values were used instead of median because some phenotypic marker’s expression was rare. Additionally, to improve signal quality, for expression markers AS, CD47, DAT, GAD65, Synaptobrevin2, vGLUT, and VMAT2 in subpopulation B1 and B2, which overlapped with postsynaptic events (**Fig. S5D** to **F**), their values were subtracted by mean values from the corresponding postsynaptic subpopulations as these markers were known to be highly enriched in pre-synapses. The preliminary robustness and sanity check of the resulting clusters compared to existing literature were detailed in **Supplementary Method** (*57, 58*).

### Machine learning model evaluation and interpretation

Similar to single cell analyses in AD and other diseases (*59, 60*), conventional machine learning models were used for prediction and identification of important features. The input to the models was the mean expression matrix obtained after clustering. For all of the classification and regression tasks in this study, five ML algorithms were compared, including LASSO, ridge, elastic net (EN), random forest, extreme gradient boosting (XGBoost), and support vector machine (SVM). Only the best performing model was reported. All of the models were run on Python using either sklearn or xgboost packages with default hyper parameters as optimizing these models to obtain the best possible prediction results was not the goal of this study. A leave-one-out cross validation method was employed to evaluate the performance of each of the models. The aggregated test prediction values were used for evaluation. The area under the receiver operating curve (AUC) and Wilcoxon’s P value were used as criteria.

### Multiplexed ion beam imaging (MIBI) for identification of subpopulation characteristics suggested by SynTOF

ADNC and Control hippocampus samples from different brain autopsies than those used for SynTOF were sectioned at 4 mm thickness from archival FFPE tissue blocks and mounted on salinized gold slides for MIBI-TOF. MIBI-TOF antibody conjugation and staining exactly followed previously published protocols (*61, 62*). Acquisition of spectral images was performed using a modified MIBI-TOF (Oregon Physics & IONPATH) with a hyperion ion source (*32*). Images were preprocessed as described previously (*32*).

To assign meaningful phenotypes to each pixel of the image, we applied a pixel clustering approach. Pre-processed (background subtracted, de-noised) MIBI images were first Gaussian blurred using a standard deviation of 2 for the Gaussian kernel. Pixels were normalized by their total expression, such that the total expression of each pixel was equal to 1. A 99.9% normalization was applied for each marker. Pixels were clustered into 100 clusters using FlowSOM based on the expression of the markers. The average expression of each of the 100 pixel clusters was found and the z-score for each marker across the 100 pixel clusters was computed. All z-scores were capped at 3, such that the maximum z-score was 3. Using these z-scored expression values, the 100 pixel clusters were clustered using consensus hierarchical clustering using Euclidean distance into 20 metaclusters. In our experience, for a set of 16 clustering markers, 20 metaclusters are able to capture the phenotypic diversity well. Upon inspection of the expression heatmap of the 20 metaclusters, we observed discrete phenotypes for each metacluster. These metaclusters were mapped back to the original images to generate overlay images colored by pixel metacluster. Each pixel was therefore assigned to an interpretable phenotypic cluster.

We then applied a sliding window approach to assess the co-localization of proteins within each image. We varied the size of the sliding window, using 3×3 pixels windows (0.9 μm^2^) with a stride of 2 pixels, 6×6 (0.9 μm^2^) windows with a stride of 3 pixels, and 9×9 pixels (2.7 μm^2^) windows with a stride of 3 pixels. Results were consistent across different window sizes. For each window, we assessed marker positivity by counting the number of pixels of each pixel cluster within each window. For example, if a window contained a pixel belonging to pixel cluster 1, which was defined by the Tau expression, the window was called as Tau+. For each window, we assessed marker positivity for CD56, vGLUT, ApoE, GAD65, and tau. We varied the cut-off for calling positivity and results were consistent across different cut-offs. By using such an approach, we identified which markers were expressed in each window. By assessing combinations of markers within each window, we could investigate marker co-localization.

Furthermore, we assessed the mean expression of CD47 and PHF-tau within each window. The MIBI images were transformed by multiplying by 100 and arcsinh transformed with a cofactor of 5, then rescaled on a 0 to 5 scale. The images were then normalized by average histone H3 (HH3) expression to mitigate batch effects between MIBI images, since average HH3 expression is consistent across images (*32*). The average expression of CD47 and PHF-tau were then calculated using these normalized values.

### Statistical Methods

Prism software and R programming language were used to conduct statistical analysis. All single-event/single-pixel level analyses and the Control’s subpopulation frequency are presented in bar graphs with means ± SEM and were analyzed using two-way ANOVA or Wilcoxon tests with Tukey’s correction if needed for multiple group comparisons. Differences were considered significant at P<0.001 for single synapse events and <0.01 for Control’s subpopulation frequency. The significant mean value expressions were determined from the false discovery rate (FDR) adjusted P value of the Spearman’s correlations between the features and the diagnoses. Differences were considered significant if Q<0.05. Selected significant features were also visualized in boxplots with median and quantiles values with Wilcoxon P values.

## Supplementary Materials

### Materials and Methods

Fig. S1. Comparison of % positive events between human and nonhuman primate synaptosome obtained from an ideal laboratory conditions confirmed synaptosome integrity.

Fig. S2. Comparison of the SynTOF % positive events to known ultrastructural studies indicate good agreement.

Fig. S3. Heatmaps of phenotypic markers’ mean expressions in each brain region and from a leave-one-out clustering robustness analysis.

Fig. S4. A 2-D t-SNE projections colored by phenotypic intensity confirm phenotypic definitions of each subpopulation.

Fig. S5. The pre- and postsynaptic subpopulation frequency across all regions and diagnoses show little difference between diagnoses and that postsynaptic events are mostly in B subpopulations.

Fig. S6. Expression of markers with complete separation between Control and AD groups in almost all subpopulations.

Fig. S7. The tabulated list of all significant features between AD vs. Control and non-resilient AD vs. Control.

Fig. S8. Similar MIBI analyses to Fig. 5 but with different sliding window sizes resulted in the same conclusions.

Fig. S9. DAT expression in all subpopulations of the three regions conforms with traditional knowledge.

Table S1. Characteristics of Human Samples.

Table S2. SynTOF antibody panel.

Table S3. Features (brain region, marker expression, subpopulation) that exhibited complete separation between Control vs. ADNC or Control vs. LBD.

## Acknowledgments

The authors gratefully acknowledge Prof. Katrin I. Andreasson for providing PS/APP mice.

## Author contributions

Conceptualization: TJM, NA, SCB, CRG, TP

Resources: NP, ES, CL, CAS, TCR, SC, KLP, CDK

Methodology: CRG, R Fernandez, P, ALC, MB, R Fallahzadeh, KV, TJM, NA, SCB, MA

Investigation: CRG, R Fernandez, KV

Software: TP, CCL, ALC, MB, R Fallahzadeh

Visualization: TP, TJM, CLL, KV, KSP

Funding acquisition: TJM, NA, CAS, SC, TCR, KSP, EJF

Project administration: TJM, TP

Supervision: TJM

Writing – original draft: TJM, TP, CRG, CCL, KV

Writing – review & editing: NA, SCB, MA, CDK, KLP, EJF, KSM, SC, CAS, ALC, MB, R Fallahzadeh, NP

## Data and materials availability

The single synapse data are available upon request to the corresponding author. The code for clustering and subsequent analysis of the data are available at https://github.com/tpjoe/SynTOF2021.

## References and Notes

1. S. W. Scheff, D. A. Price, Synaptic pathology in Alzheimer’s disease: a review of ultrastructural studies, Neurobiol. Aging 24, 1029–1046 (2003).

2. R. D. Terry, E. Masliah, D. P. Salmon, N. Butters, R. DeTeresa, R. Hill, L. A. Hansen, R. Katzman, Physical basis of cognitive alterations in Alzheimer’s disease: synapse loss is the major correlate of cognitive impairment, Ann. Neurol. 30, 572–580 (1991).

3. E. Masliah, R. D. Terry, M. Alford, R. DeTeresa, L. A. Hansen, Cortical and subcortical patterns of synaptophysinlike immunoreactivity in Alzheimer’s disease, Am. J. Pathol. 138, 235–246 (1991).

4. B. T. Hyman, G. W. Van Hoesen, L. J. Kromer, A. R. Damasio, Perforant pathway changes and the memory impairment of Alzheimer’s disease, Ann. Neurol. 20, 472–481 (1986).

5. S. T. DeKosky, S. W. Scheff, Synapse loss in frontal cortex biopsies in Alzheimer’s disease: correlation with cognitive severity, Ann. Neurol. 27, 457–464 (1990).

6. K. H. Gylys, T. Bilousova, Flow Cytometry Analysis and Quantitative Characterization of Tau in Synaptosomes from Alzheimer’s Disease Brains, Methods Mol. Biol. 1523, 273–284 (2017).

7. B. C. Yoo, N. Cairns, M. Fountoulakis, G. Lubec, Synaptosomal proteins, beta-soluble N-ethylmaleimide-sensitive factor attachment protein (beta-SNAP), gamma-SNAP and synaptotagmin I in brain of patients with Down syndrome and Alzheimer’s disease, Dement. Geriatr. Cogn. Disord. 12, 219–225 (2001).

8. T. S. Wijasa, M. Sylvester, N. Brocke-Ahmadinejad, S. Schwartz, F. Santarelli, V. Gieselmann, T. Klockgether, F. Brosseron, M. T. Heneka, Quantitative proteomics of synaptosome S-nitrosylation in Alzheimer’s disease, J. Neurochem. 152, 710–726 (2020).

9. B. G. Perez-Nievas, T. D. Stein, H.-C. Tai, O. Dols-Icardo, T. C. Scotton, I. Barroeta-Espar, L. Fernandez-Carballo, E. L. de Munain, J. Perez, M. Marquie, A. Serrano-Pozo, M. P. Frosch, V. Lowe, J. E. Parisi, R. C. Petersen, M. D. Ikonomovic, O. L. López, W. Klunk, B. T. Hyman, T. Gómez-Isla, Dissecting phenotypic traits linked to human resilience to Alzheimer’s pathology, Brain 136, 2510–2526 (2013).

10. C. M. Henstridge, R. J. Jackson, J. M. Kim, A. G. Herrmann, A. K. Wright, S. E. Harris, M. E. Bastin, J. M. Starr, J. Wardlaw, T. H. Gillingwater, C. Smith, C.-A. McKenzie, S. R. Cox, I. J. Deary, T. L. Spires-Jones, Post-mortem brain analyses of the Lothian Birth Cohort 1936: extending lifetime cognitive and brain phenotyping to the level of the synapse, Acta Neuropathol Commun 3, 53 (2015).

11. M. Colom-Cadena, J. Pegueroles, A. G. Herrmann, C. M. Henstridge, L. Muñoz, M. Querol-Vilaseca, C. S. Martín-Paniello, J. Luque-Cabecerans, J. Clarimon, O. Belbin, R. Núñez-Llaves, R. Blesa, C. Smith, C.-A. McKenzie, M. P. Frosch, A. Roe, J. Fortea, J. Andilla, P. Loza-Alvarez, E. Gelpi, B. T. Hyman, T. L. Spires-Jones, A. Lleó, Synaptic phosphorylated α-synuclein in dementia with Lewy bodies, Brain 140, 3204–3214 (2017).

12. H.-C. Tai, B. Y. Wang, A. Serrano-Pozo, M. P. Frosch, T. L. Spires-Jones, B. T. Hyman, Frequent and symmetric deposition of misfolded tau oligomers within presynaptic and postsynaptic terminals in Alzheimer’s disease, Acta Neuropathol Commun 2, 146 (2014).

13. C. R. Gajera, R. Fernandez, N. Postupna, K. S. Montine, E. J. Fox, D. Tebaykin, M. Angelo, S. C. Bendall, C. D. Keene, T. J. Montine, Mass synaptometry: High-dimensional multi parametric assay for single synapses, J. Neurosci. Methods 312, 73–83 (2019).

14. C. R. Gajera, R. Fernandez, K. S. Montine, E. J. Fox, D. Mrdjen, N. O. Postupna, C. D. Keene, S. C. Bendall, T. J. Montine, Mass-tag barcoding for multiplexed analysis of human synaptosomes and other anuclear events, Cytometry A (2021), doi:10.1002/cyto.a.24340.

15. B. T. Hyman, C. H. Phelps, T. G. Beach, E. H. Bigio, N. J. Cairns, M. C. Carrillo, D. W. Dickson, C. Duyckaerts, M. P. Frosch, E. Masliah, S. S. Mirra, P. T. Nelson, J. A. Schneider, D. R. Thal, B. Thies, J. Q. Trojanowski, H. V. Vinters, T. J. Montine, National Institute on Aging-Alzheimer’s Association guidelines for the neuropathologic assessment of Alzheimer’s disease, Alzheimers. Dement. 8, 1–13 (2012).

16. T. J. Montine, C. H. Phelps, T. G. Beach, E. H. Bigio, N. J. Cairns, D. W. Dickson, C. Duyckaerts, M. P. Frosch, E. Masliah, S. S. Mirra, P. T. Nelson, J. A. Schneider, D. R. Thal, J. Q. Trojanowski, H. V. Vinters, B. T. Hyman, National Institute on Aging, Alzheimer’s Association, National Institute on Aging-Alzheimer’s Association guidelines for the neuropathologic assessment of Alzheimer’s disease: a practical approach, Acta Neuropathol. 123, 1–11 (2012).

17. T. J. Montine, S. A. Bukhari, L. R. White, Cognitive Impairment in Older Adults and Therapeutic Strategies, Pharmacol. Rev. 73, 152–162 (2021).

18. L. Holcomb, M. N. Gordon, E. McGowan, X. Yu, S. Benkovic, P. Jantzen, K. Wright, I. Saad, R. Mueller, D. Morgan, S. Sanders, C. Zehr, K. O’Campo, J. Hardy, C. M. Prada, C. Eckman, S. Younkin, K. Hsiao, K. Duff, Accelerated Alzheimer-type phenotype in transgenic mice carrying both mutant amyloid precursor protein and presenilin 1 transgenes, Nat. Med. 4, 97–100 (1998).

19. N. Postupna, C. S. Latimer, E. B. Larson, E. Sherfield, J. Paladin, C. A. Shively, M. J. Jorgensen, R. N. Andrews, J. R. Kaplan, P. K. Crane, K. S. Montine, S. Craft, C. D. Keene, T. J. Montine, Human Striatal Dopaminergic and Regional Serotonergic Synaptic Degeneration with Lewy Body Disease and Inheritance of APOE ε4, Am. J. Pathol. 187, 884–895 (2017).

20. P. Somogyi, G. Tamás, R. Lujan, E. H. Buhl, Salient features of synaptic organisation in the cerebral cortex1Published on the World Wide Web on 3 March 1998.1 Brain Research Reviews 26, 113–135 (1998).

21. J. P. Bolam, J. Paul Bolam, E. K. Pissadaki, Living on the edge with too many mouths to feed: Why dopamine neurons dieMovement Disorders 27, 1478–1483 (2012).

22. J. F. Smiley, P. S. Goldman-Rakic, Heterogeneous Targets of Dopamine Synapses in Monkey Prefrontal Cortex Demonstrated by Serial Section Electron Microscopy: A Laminar Analysis Using the Silver-enhanced Diaminobenzidine Sulfide (SEDS) Immunolabeling Technique Cerebral Cortex 3, 223–238 (1993).

23. C. J. Lacey, J. Boyes, O. Gerlach, L. Chen, P. J. Magill, J. P. Bolam, GABA(B) receptors at glutamatergic synapses in the rat striatum, Neuroscience 136, 1083–1095 (2005).

24. V. J. Roberts, S. L. Barth, H. Meunier, W. Vale, Hybridization histochemical and immunohistochemical localization of inhibin/activin subunits and messenger ribonucleic acids in the rat brain The Journal of Comparative Neurology 364, 473–493 (1996).

25. G. W. Miller, J. D. Erickson, J. T. Perez, S. N. Penland, D. C. Mash, D. B. Rye, A. I. Levey, Immunochemical Analysis of Vesicular Monoamine Transporter (VMAT2) Protein in Parkinson’s Disease Experimental Neurology 156, 138–148 (1999).

26. J. Booij, G. Tissingh, G. J. Boer, J. D. Speelman, J. C. Stoof, A. G. Janssen, E. C. Wolters, E. A. van Royen, [123I]FP-CIT SPECT shows a pronounced decline of striatal dopamine transporter labelling in early and advanced Parkinson’s disease, J. Neurol. Neurosurg. Psychiatry 62, 133–140 (1997).

27. H. Sasaguri, P. Nilsson, S. Hashimoto, K. Nagata, T. Saito, B. De Strooper, J. Hardy, R. Vassar, B. Winblad, T. C. Saido, APP mouse models for Alzheimer’s disease preclinical studies, EMBO J. 36, 2473–2487 (2017).

28. L. Mucke, D. J. Selkoe, Neurotoxicity of amyloid β-protein: synaptic and network dysfunction, Cold Spring Harb. Perspect. Med. 2, a006338 (2012).

29. N. Aghaeepour, E. F. Simonds, D. J. H. F. Knapp, R. V. Bruggner, K. Sachs, A. Culos, P. F. Gherardini, N. Samusik, G. K. Fragiadakis, S. C. Bendall, B. Gaudilliere, M. S. Angst, C. J. Eaves, W. A. Weiss, W. J. Fantl, G. P. Nolan, GateFinder: projection-based gating strategy optimization for flow and mass cytometry, Bioinformatics 34, 4131–4133 (2018).

30. I. G. McKeith, B. F. Boeve, D. W. Dickson, G. Halliday, J.-P. Taylor, D. Weintraub, D. Aarsland, J. Galvin, J. Attems, C. G. Ballard, A. Bayston, T. G. Beach, F. Blanc, N. Bohnen, L. Bonanni, J. Bras, P. Brundin, D. Burn, A. Chen-Plotkin, J. E. Duda, O. El-Agnaf, H. Feldman, T. J. Ferman, D. Ffytche, H. Fujishiro, D. Galasko, J. G. Goldman, S. N. Gomperts, N. R. Graff-Radford, L. S. Honig, A. Iranzo, K. Kantarci, D. Kaufer, W. Kukull, V. M. Y. Lee, J. B. Leverenz, S. Lewis, C. Lippa, A. Lunde, M. Masellis, E. Masliah, P. McLean, B. Mollenhauer, T. J. Montine, E. Moreno, E. Mori, M. Murray, J. T. O’Brien, S. Orimo, R. B. Postuma, S. Ramaswamy, O. A. Ross, D. P. Salmon, A. Singleton, A. Taylor, A. Thomas, P. Tiraboschi, J. B. Toledo, J. Q. Trojanowski, D. Tsuang, Z. Walker, M. Yamada, K. Kosaka, Diagnosis and management of dementia with Lewy bodies: Fourth consensus report of the DLB Consortium, Neurology 89, 88– 100 (2017).

31. M. Ballmaier, J. T. O’Brien, E. J. Burton, P. M. Thompson, D. E. Rex, K. L. Narr, I. G. McKeith, H. DeLuca, A. W. Toga, Comparing gray matter loss profiles between dementia with Lewy bodies and Alzheimer’s disease using cortical pattern matching: diagnosis and gender effects, Neuroimage 23, 325–335 (2004).

32. L. Keren, M. Bosse, S. Thompson, T. Risom, K. Vijayaragavan, E. McCaffrey, D. Marquez, R. Angoshtari, N. F. Greenwald, H. Fienberg, J. Wang, N. Kambham, D. Kirkwood, G. Nolan, T. J. Montine, S. J. Galli, R. West, S. C. Bendall, M. Angelo, MIBI-TOF: A multiplexed imaging platform relates cellular phenotypes and tissue structure, Sci Adv 5, eaax5851 (2019).

33. V. Rangaraju, N. Calloway, T. A. Ryan, Activity-Driven Local ATP Synthesis Is Required for Synaptic Function Cell 156, 825–835 (2014).

34. M. Gratuze, D. M. Holtzman, Targeting pre-synaptic tau accumulation: a new strategy to counteract tau-mediated synaptic loss and memory deficits Neuron 109, 741–743 (2021).

35. J. Choi, M. C. Sullards, J. A. Olzmann, H. D. Rees, S. T. Weintraub, D. E. Bostwick, M. Gearing, A. I. Levey, L.-S. Chin, L. Li, Oxidative damage of DJ-1 is linked to sporadic Parkinson and Alzheimer diseases, J. Biol. Chem. 281, 10816–10824 (2006).

36. K. Solti, W.-L. Kuan, B. Fórizs, G. Kustos, J. Mihály, Z. Varga, B. Herberth, É. Moravcsik, R. Kiss, M. Kárpáti, A. Mikes, Y. Zhao, T. Imre, J.-C. Rochet, F. Aigbirhio, C. H. Williams-Gray, R. A. Barker, G. Tóth, DJ-1 can form β-sheet structured aggregates that co-localize with pathological amyloid deposits, Neurobiol. Dis. 134, 104629 (2020).

37. D. J. Moore, L. Zhang, J. Troncoso, M. K. Lee, N. Hattori, Y. Mizuno, T. M. Dawson, V. L. Dawson, Association of DJ-1 and parkin mediated by pathogenic DJ-1 mutations and oxidative stress, Hum. Mol. Genet. 14, 71–84 (2005).

38. P. Heutink, PINK-1 and DJ-1--new genes for autosomal recessive Parkinson’s disease, J. Neural Transm. Suppl., 215–219 (2006).

39. T. J. Montine, K. S. Montine, W. McMahan, W. R. Markesbery, J. F. Quinn, J. D. Morrow, F2-isoprostanes in Alzheimer and other neurodegenerative diseases, Antioxid. Redox Signal. 7, 269–275 (2005).

40. H. Ohnishi, Y. Kaneko, H. Okazawa, M. Miyashita, R. Sato, A. Hayashi, K. Tada, S. Nagata, M. Takahashi, T. Matozaki, Differential localization of Src homology 2 domain-containing protein tyrosine phosphatase substrate-1 and CD47 and its molecular mechanisms in cultured hippocampal neurons, J. Neurosci. 25, 2702–2711 (2005).

41. A. B. Toth, A. Terauchi, L. Y. Zhang, E. M. Johnson-Venkatesh, D. J. Larsen, M. A. Sutton, H. Umemori, Synapse maturation by activity-dependent ectodomain shedding of SIRPα, Nat. Neurosci. 16, 1417–1425 (2013).

42. T. W. Miller, J. S. Isenberg, H. B. Shih, Y. Wang, D. D. Roberts, Amyloid-β inhibits No-cGMP signaling in a CD36-and CD47-dependent manner, PLoS One 5, e15686 (2010).

43. M. H. Han, D. H. Lundgren, S. Jaiswal, M. Chao, K. L. Graham, C. S. Garris, R. C. Axtell, P. P. Ho, C. B. Lock, J. I. Woodard, S. E. Brownell, M. Zoudilova, J. F. V. Hunt, S. E. Baranzini, E. C. Butcher, C. S. Raine, R. A. Sobel, D. K. Han, I. Weissman, L. Steinman, Janus-like opposing roles of CD47 in autoimmune brain inflammation in humans and mice, J. Exp. Med. 209, 1325–1334 (2012).

44. E. K. Lehrman, D. K. Wilton, E. Y. Litvina, C. A. Welsh, S. T. Chang, A. Frouin, A. J. Walker, M. D. Heller, H. Umemori, C. Chen, B. Stevens, CD47 Protects Synapses from Excess Microglia-Mediated Pruning during Development, Neuron 100, 120–134.e6 (2018).

45. T. Bartels, S. De Schepper, S. Hong, Microglia modulate neurodegeneration in Alzheimer’s and Parkinson’s diseases Science 370, 66–69 (2020).

46. M. P. Chao, C. H. Takimoto, D. D. Feng, K. McKenna, P. Gip, J. Liu, J.-P. Volkmer, I. L. Weissman, R. Majeti, Therapeutic Targeting of the Macrophage Immune Checkpoint CD47 in Myeloid Malignancies, Front. Oncol. 9, 1380 (2019).

47. J. D. Ulrich, T. K. Ulland, T. E. Mahan, S. Nyström, K. P. Nilsson, W. M. Song, Y. Zhou, M. Reinartz, S. Choi, H. Jiang, F. R. Stewart, E. Anderson, Y. Wang, M. Colonna, D. M. Holtzman, ApoE facilitates the microglial response to amyloid plaque pathology, J. Exp. Med. 215, 1047–1058 (2018).

48. Y. Shi, K. Yamada, S. A. Liddelow, S. T. Smith, L. Zhao, W. Luo, R. M. Tsai, S. Spina, L. T. Grinberg, J. C. Rojas, G. Gallardo, K. Wang, J. Roh, G. Robinson, M. B. Finn, H. Jiang, P. M. Sullivan, C. Baufeld, M. W. Wood, C. Sutphen, L. McCue, C. Xiong, J. L. Del-Aguila, J. C. Morris, C. Cruchaga, Alzheimer’s Disease Neuroimaging Initiative, A. L. Fagan, B. L. Miller, A. L. Boxer, W. W. Seeley, O. Butovsky, B. A. Barres, S. M. Paul, D. M. Holtzman, ApoE4 markedly exacerbates tau-mediated neurodegeneration in a mouse model of tauopathy, Nature 549, 523–527 (2017).

49. T.-P. V. Huynh, F. Liao, C. M. Francis, G. O. Robinson, J. R. Serrano, H. Jiang, J. Roh, M. B. Finn, P. M. Sullivan, T. J. Esparza, F. R. Stewart, T. E. Mahan, J. D. Ulrich, T. Cole, D. M. Holtzman, Age-Dependent Effects of apoE Reduction Using Antisense Oligonucleotides in a Model of β-amyloidosis, Neuron 96, 1013–1023.e4 (2017).

50. G. M. McKhann, D. S. Knopman, H. Chertkow, B. T. Hyman, C. R. Jack Jr, C. H. Kawas, W. E. Klunk, W. J. Koroshetz, J. J. Manly, R. Mayeux, R. C. Mohs, J. C. Morris, M. N. Rossor, P. Scheltens, M. C. Carrillo, B. Thies, S. Weintraub, C. H. Phelps, The diagnosis of dementia due to Alzheimer’s disease: recommendations from the National Institute on Aging-Alzheimer’s Association workgroups on diagnostic guidelines for Alzheimer’s disease, Alzheimers. Dement. 7, 263–269 (2011).

51. N. Stanley, I. A. Stelzer, A. S. Tsai, R. Fallahzadeh, E. Ganio, M. Becker, T. Phongpreecha, H. Nassar, S. Ghaemi, I. Maric, A. Culos, A. L. Chang, M. Xenochristou, Han, C. Espinosa, K. Rumer, L. Peterson, F. Verdonk, D. Gaudilliere, E. Tsai, D. Feyaerts, J. Einhaus, K. Ando, R. J. Wong, G. Obermoser, G. M. Shaw, D. K. Stevenson, M. S. Angst, B. Gaudilliere, N. Aghaeepour, VoPo leverages cellular heterogeneity for predictive modeling of single-cell data, Nat. Commun. 11, 3738 (2020).

52. S. Van Gassen, B. Callebaut, M. J. Van Helden, B. N. Lambrecht, P. Demeester, T. Dhaene, Y. Saeys, FlowSOM: Using self-organizing maps for visualization and interpretation of cytometry data, Cytometry A 87, 636–645 (2015).

53. L. M. Weber, M. D. Robinson, Comparison of clustering methods for high-dimensional single-cell flow and mass cytometry data, Cytometry A 89, 1084–1096 (2016).

54. X. Liu, W. Song, B. Y. Wong, T. Zhang, S. Yu, G. N. Lin, X. Ding, A comparison framework and guideline of clustering methods for mass cytometry data, Genome Biol. 20, 297 (2019).

55. X. Guo, L. Gao, X. Liu, J. Yin, Improved Deep Embedded Clustering with Local Structure Preservation Proceedings of the Twenty-Sixth International Joint Conference on Artificial Intelligence (2017), doi:10.24963/ijcai.2017/243.

56. K. Hornik, A CLUE for CLUster Ensembles Journal of Statistical Software 14 (2005), doi:10.18637/jss.v014.i12.

57. K. H. Gylys, J. A. Fein, F. Yang, D. J. Wiley, C. A. Miller, G. M. Cole, Synaptic changes in Alzheimer’s disease: increased amyloid-beta and gliosis in surviving terminals is accompanied by decreased PSD-95 fluorescence, Am. J. Pathol. 165, 1809–1817 (2004).

58. K. H. Gylys, J. A. Fein, G. M. Cole, Quantitative characterization of crude synaptosomal fraction (P-2) components by flow cytometry, J. Neurosci. Res. 61, 186–192 (2000).

59. A. Culos, A. S. Tsai, N. Stanley, M. Becker, M. S. Ghaemi, D. R. McIlwain, R. Fallahzadeh, A. Tanada, H. Nassar, C. Espinosa, M. Xenochristou, E. Ganio, L. Peterson, X. Han, I. A. Stelzer, K. Ando, D. Gaudilliere, T. Phongpreecha, I. Marić, A. L. Chang, G. M. Shaw, D. K. Stevenson, S. Bendall, K. L. Davis, W. Fantl, G. P. Nolan, T. Hastie, R. Tibshirani, M. S. Angst, B. Gaudilliere, N. Aghaeepour, Integration of mechanistic immunological knowledge into a machine learning pipeline improves predictions, Nat Mach Intell 2, 619–628 (2020).

60. T. Phongpreecha, R. Fernandez, D. Mrdjen, A. Culos, C. R. Gajera, A. M. Wawro, N. Stanley, B. Gaudilliere, K. L. Poston, N. Aghaeepour, T. J. Montine, Single-cell peripheral immunoprofiling of Alzheimer’s and Parkinson’s diseases, Sci Adv 6 (2020), doi:10.1126/sciadv.abd5575.

61. M. M. B. Bosse, S. C. Bendall, M.R. Angelo, MIBI staining V.2. dx.doi.org/10.17504/protocols.io.bt9tnr6n (2021).

62. M. M. B. Bosse, C. Camacho, S. C. Bendall, M.R. Angelo, MIBI and IHC solutions. dx.doi.org/10.17504/protocols.io.bhmej43e (2021).

