## Supplementary material for "Single-synapse analyses of Alzheimer’s disease implicate pathologic tau, DJ1, CD47, and ApoE": Fig. S1 - Fig. S9; Table S1 - Table S3

### Supplementary Materials and Methods

#### Preliminary robustness and sanity checks of the resulting clusters

To ensure that the resulting clusters were robust, a leave-one-out cluster prediction was performed on the Control samples and compared to the original results obtained from training on all 6 Control samples. Specifically, in each of the 6 iterations, the autoencoder was trained on 5 Control samples and predicted the cluster assignment for the held-out sample. The results are visualized in Fig. S3B where each pixel of the heatmap represents average scaled expression values from the predicted held-out samples. The similarity between this heatmap and that in Fig. 2A suggests that the resulting clusters are equivalent. The mean and standard deviation of the correlation statistics (Pearson's R) between them for each of the phenotypic markers are shown on the left of the heatmap.

Autoencoder-generated clusters were initially validated by two approaches. First, subpopulations followed strong expectations, *e.g,* all presynaptic subpopulations in caudate exhibited multiple fold higher DAT expression than in BA9 or hippocampus (Fig. S9). Second, the intentionally selected redundant presynaptic markers, CD47 and SNAP25, had highly correlated signal intensities across subpopulations (Spearman's R=0.90, 0.91, and 0.91 in BA9, caudate, and hippocampus, $P=2.2\times10^{-16}$ for all regions). Multiple groups have shown that a minority of synaptosomes retain attached astrocytic components *(13, 57, 58)*; the intentionally redundant astrocytic markers, EAAT1 and GFAP, exhibited strong correlations of intensities across multiple subpopulations in all regions (Spearman's R=0.74, 0.62, and 0.68 in BA9, caudate, and hippocampus, $P<2.2\times10^{-16}$ for all regions).

**Supplementary Figures**

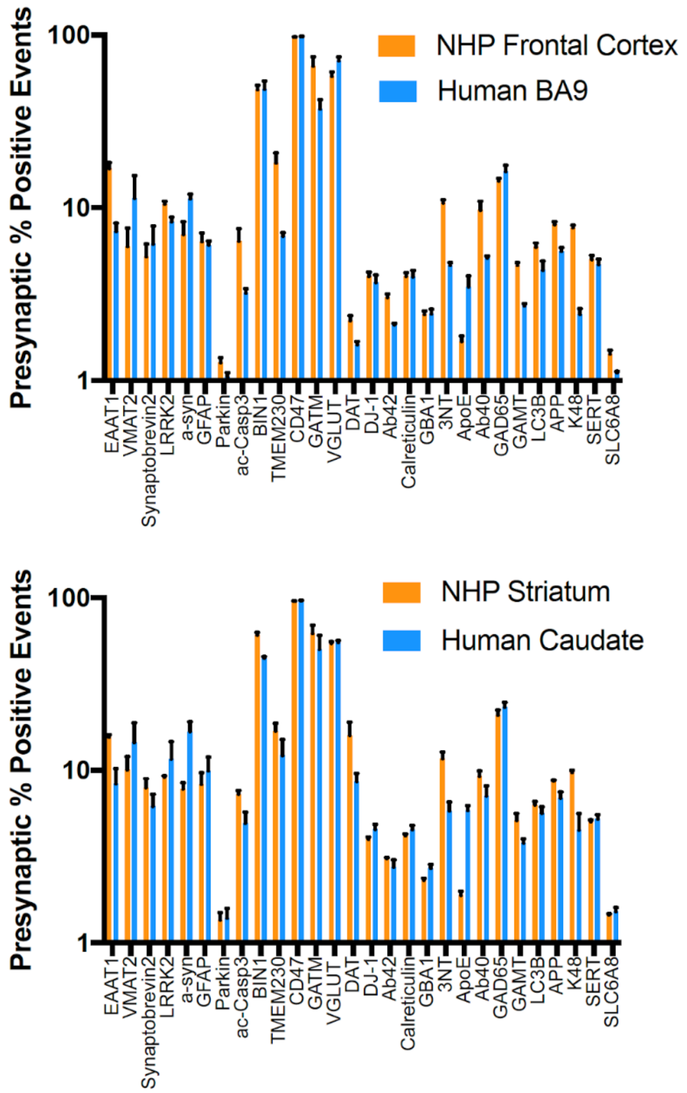


**Fig. S1**. **Comparison of % positive events between human and nonhuman primate synaptosome obtained from an ideal laboratory conditions confirmed synaptosome integrity.** The molecular integrity of human synaptosome preparations from Control BA9 and caudate was tested by comparison with four adult (11.5±0.3) female nonhuman primate (NHP) synaptosome preparations from midfrontal lobe and striatum that were obtained under ideal research conditions. Antibodies to human tau, PHF-tau, p129-α-synclein, and PrP did not cross react with NHP and were dropped. The only significant differences between Control and NHP were greater vGLUT ($P<0.01$) and less GATM ($P<0.001$) in human frontal cortex, and less BIN1 ($P<0.001$) in human striatum. There was no consistent pattern of reduced signal in human synaptosome preparations highlighting their molecular integrity when prepared as described here.

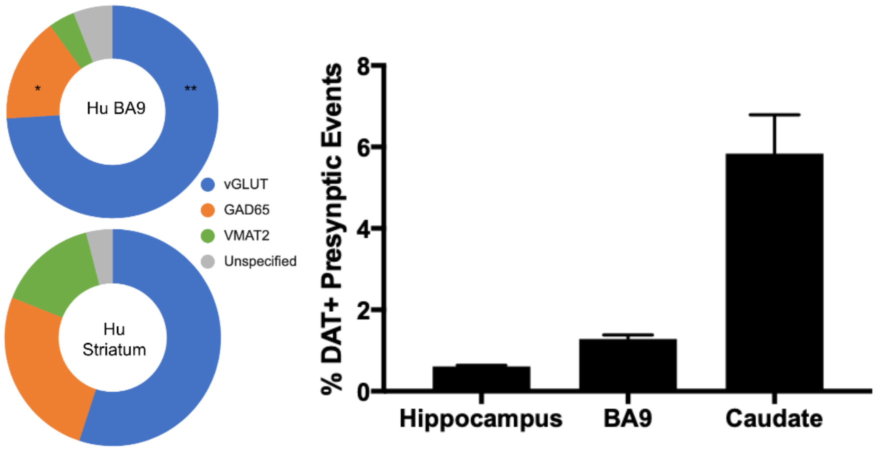


**Fig. S2. Comparison of the SynTOF % positive events to known ultrastructural studies indicate good agreement.** Ultrastructural studies have estimated the proportion of different types of synapses in primate brain. In prefrontal cortex, approximately 80% of synapses are asymmetric and 85% to 90% of these are glutamatergic, aligning well with the SynTOF estimate of 73.0±3.5% BA9 vGLUT+ synaptosomes in Controls *(20)*. The approximately 20% symmetric synapses in prefrontal cortex are mostly GABAergic, estimated at 1 in 6 or 17% of total *(20)*. The percent of BA9 GAD65+ presynaptic events by SynTOF was 16.3±1.4% in Controls. The remaining symmetric synapses (about 3% of total) and some of the non-glutamatergic asymmetric synapses are monoaminergic, both noradrenergic and dopaminergic at this site, although ultrastructural assessments of monoaminergic terminals are likely underestimates because many terminals are not well formed synapses *(21, 22)*. BA9 VMAT2+ events were 4.3±0.9% of Control total presynaptic events by SynTOF. In primate caudate nucleus, 70% to 86% of synapses are asymmetric but only two-thirds, or 47% to 58%, are glutamatergic *(23)*. Control caudate vGLUT+ presynaptic events were 55.0±1.5% of total by SynTOF, which was significantly less than in BA9 (**P<0.01). The remaining asymmetric synapses in caudate are mostly serotonergic with some dopaminergic. Symmetric synapses in the caudate nucleus range from 24% to 30% and are mostly GABAergic along with dopaminergic *(23, 24)*. Control caudate GAD65+ presynaptic events were 26.2±1.6%, by SynTOF, which was significantly more than BA9 (*P<0.05). VMAT2+ presynaptic events in Control caudate were 14.5±2.6% of total by SynTOF. The ratio of DAT to VMAT2 varies between 0.5 and 0.3 in human striatum *(25)*. DAT+/VAMT2+ presynaptic events by SynTOF averaged 0.4 in Controls. As expected, distribution of DAT+ presynaptic events by SynTOF was significantly different across the three regions in Controls (P<0.0001) with caudate approximately 5-fold greater than BA9 (P<0.001) and 10-fold greater than hippocampus *(25)* (P<0.001).

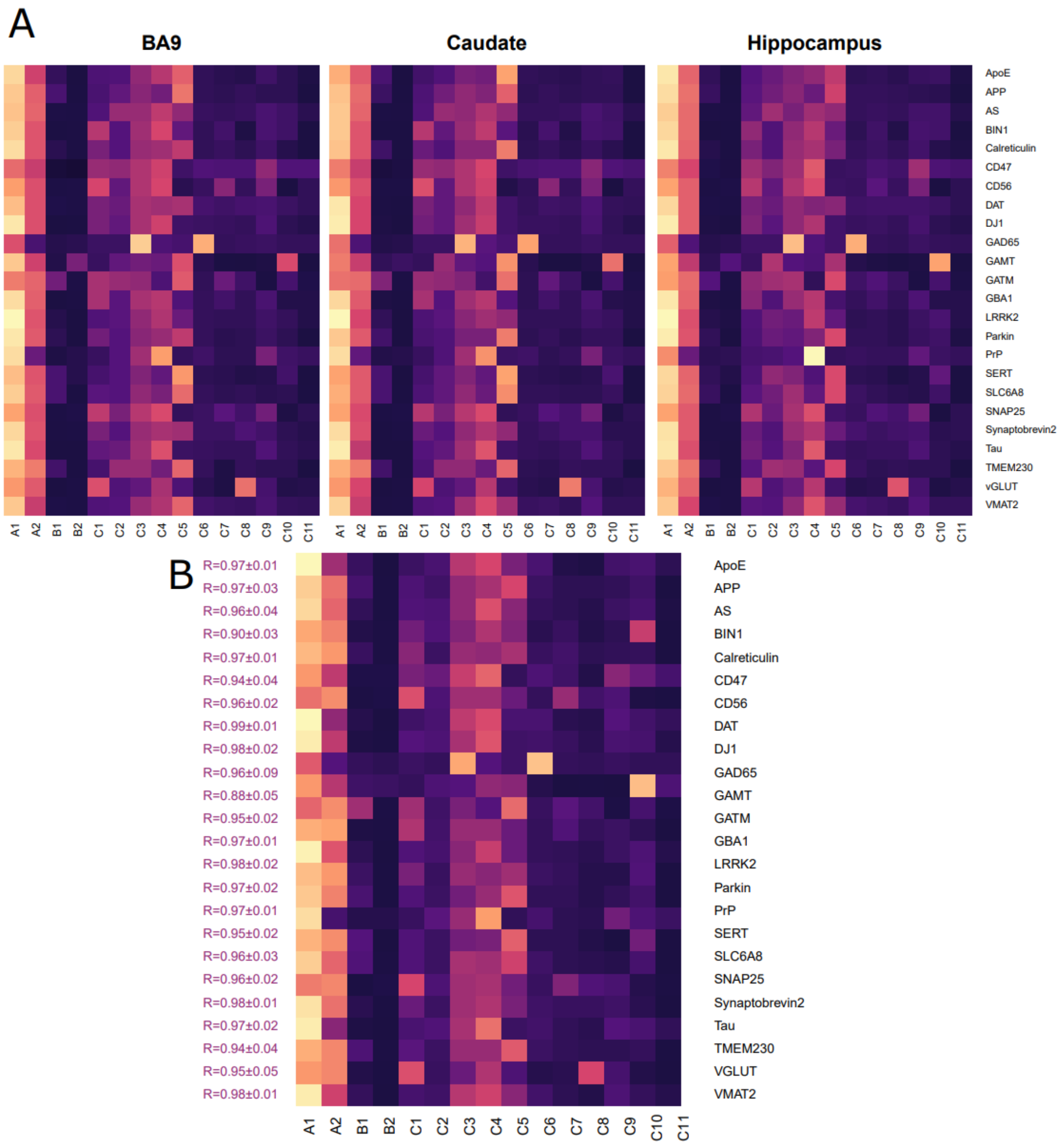


**Fig. S3. Heatmaps of phenotypic markers' mean expressions in each brain region and from a leave-one-out clustering robustness analysis.** (**A**) A heatmap showing the mean expression values from Control samples for each of the phenotypic markers across all identified subpopulations and stratified by brain region. The mean expression values were scaled across subpopulations for differential visualization purposes. These heatmaps demonstrate that the subpopulation characteristics were similar across all brain regions. (**B**) A heatmap showing the mean expression values from the average of the leave-one-out prediction of the six Control samples (all brain regions). For each of the phenotypic markers, the correlations to the mean expression values obtained from training and predicting using all six samples (as shown in Fig. 2A) were measured by Pearson's R. The high correlations between the two indicate the robustness of the clustering results. The mean expression values were scaled across subpopulations for visualization purposes.

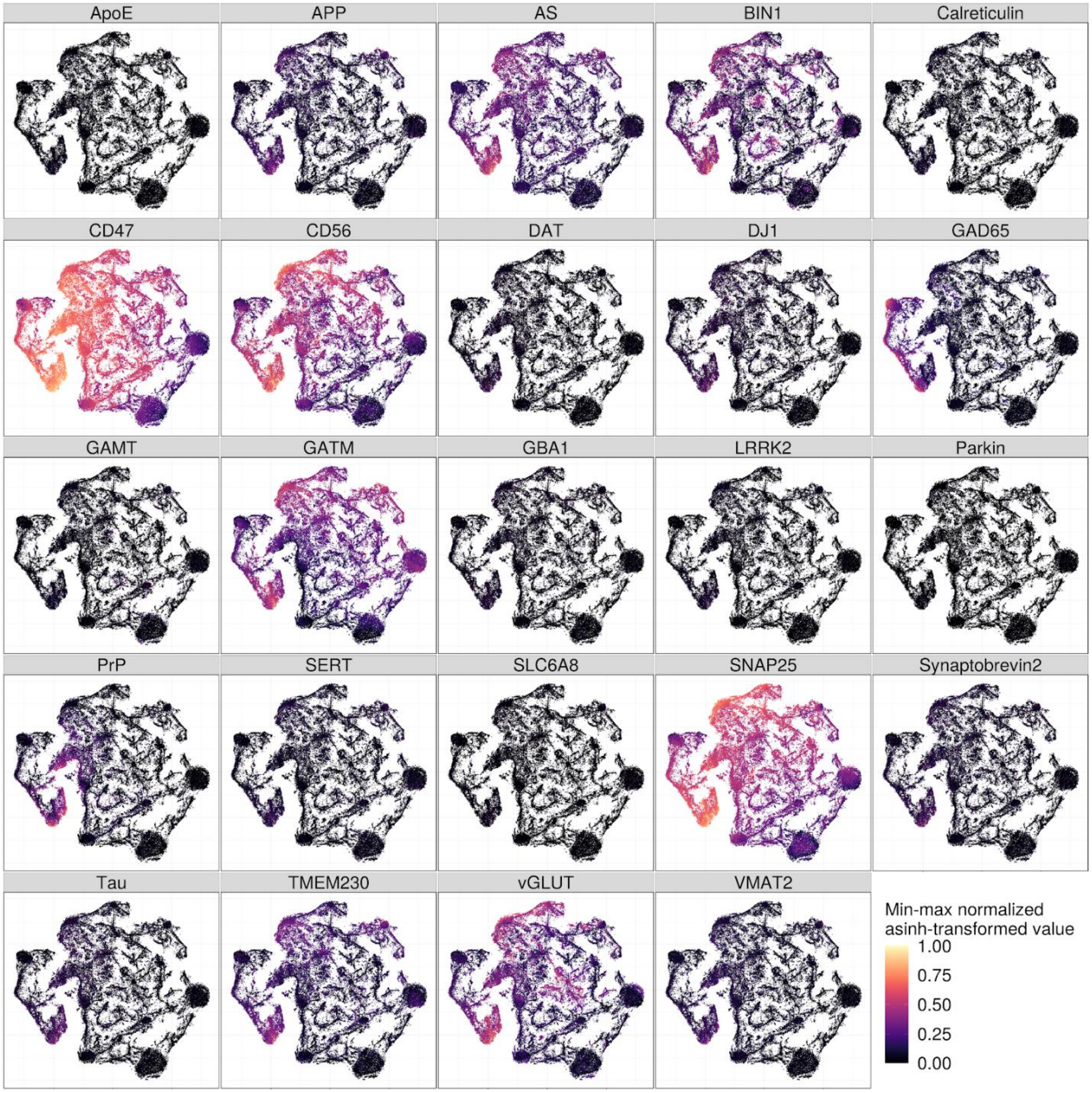


**Fig. S4**. **A 2-D t-SNE projections colored by phenotypic intensity confirm phenotypic definitions of each subpopulation.** The t-SNE plot of the sampled autoencoder's hidden representations from all brain regions of the Control samples with each event colored by the intensity of each of the defined phenotypic markers.

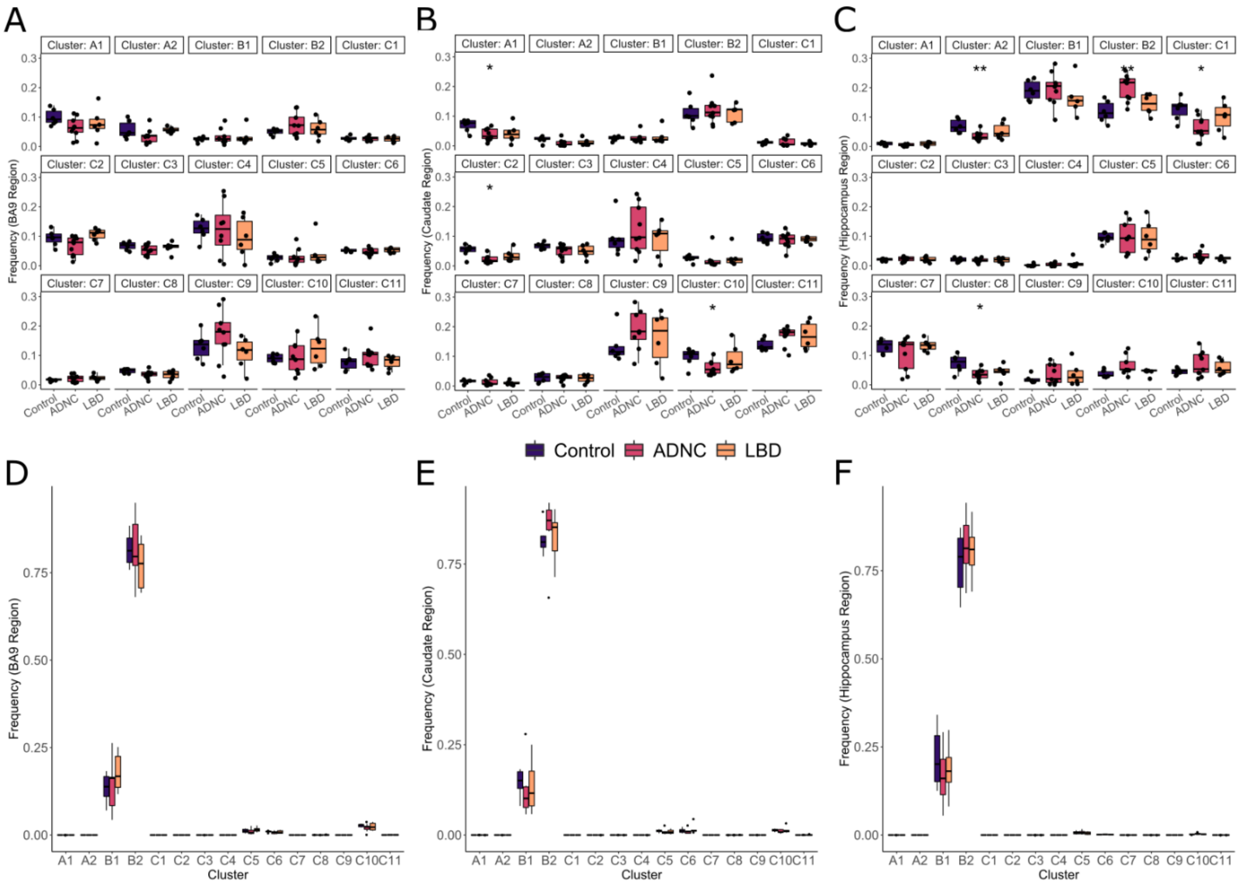


**Fig. S5. The pre- and postsynaptic subpopulation frequency across all regions and diagnoses show little difference between diagnoses and that postsynaptic events are mostly in B subpopulations.** (**A**)-(**C**) The frequency of each subpopulation of presynaptic events for Control, ADNC, and LBD in BA9, caudate, and hippocampus region, respectively. Asterisks indicate that the frequency of the disease group is significantly different (Wilcoxon) from Control. (**D**)-(**F**) The frequency of each subpopulation of postsynaptic events for Control, ADNC, and LBD in BA9, caudate, and hippocampus region, respectively. Most postsynaptic events were similar to subpopulation B.

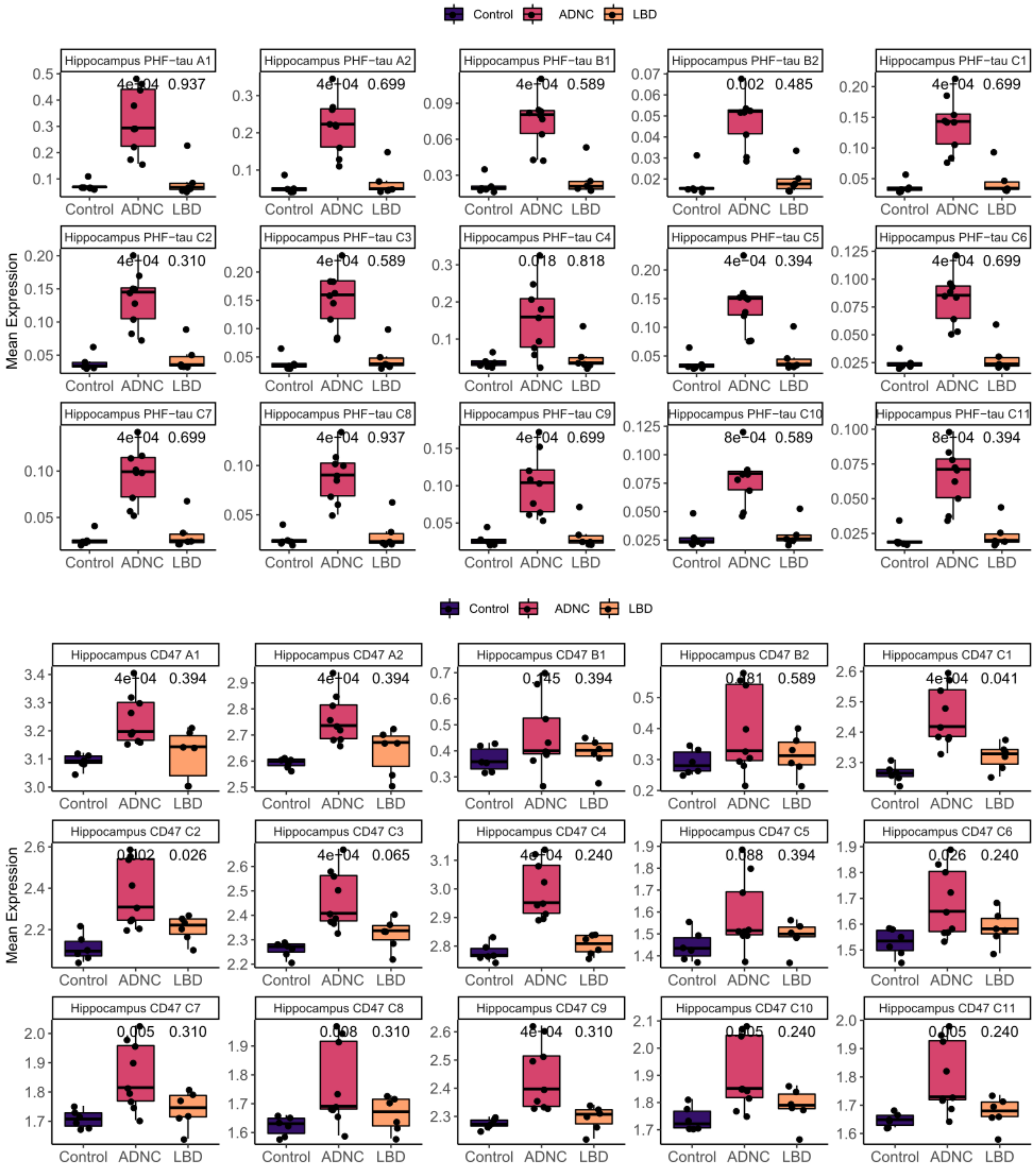


**Fig. S6**. **Expression of markers with complete separation between Control and AD groups in almost all subpopulations.** These include PHF-tau expression for almost all subpopulations in hippocampus (top) and CD47 expression for most subpopulations with higher CD47 expressions in hippocampus (bottom), including subpopulation A1 and A2, C1to C4, and C7 to C11.

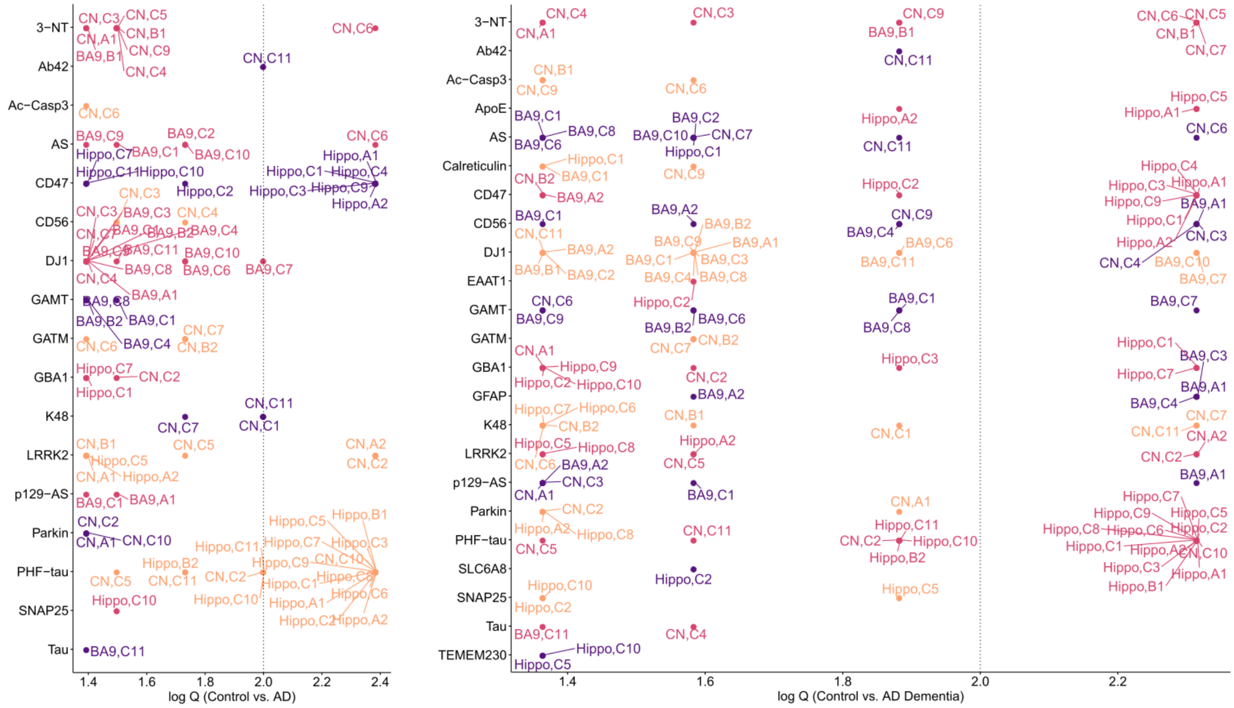


**Fig. S7**. **The tabulated list of all significant features between AD vs. Control and non-resilient AD vs. Control.** The list of all significant features (Q value < 0.05) for Control (n=6) vs. AD (n=9) (left) stratified by marker type shows prevalent and strong signals from PHF-tau and CD47 expression in multiple hippocampal subpopulations. Less strong, but still prevalent expression signals are DJ1 in multiple BA9 subpopulations. Other notable signals include GATM and K48 in caudate, and 3-NT in BA9. The right panel shows a list of all significant features (Q value < 0.05) for Control (n=6) vs. AD dementia (n=7), in which most signals from the previous comparison became stronger. Notably, ApoE was one of the few new signals (absent in the left panel) that also ranked among the strongest. Markers omitted on the y-axis were those without any significant subpopulations in any brain regions. Color is included only to facilitate visualization of each row. Note that CN and Hippo are abbreviations for caudate and hippocampus, respectively.

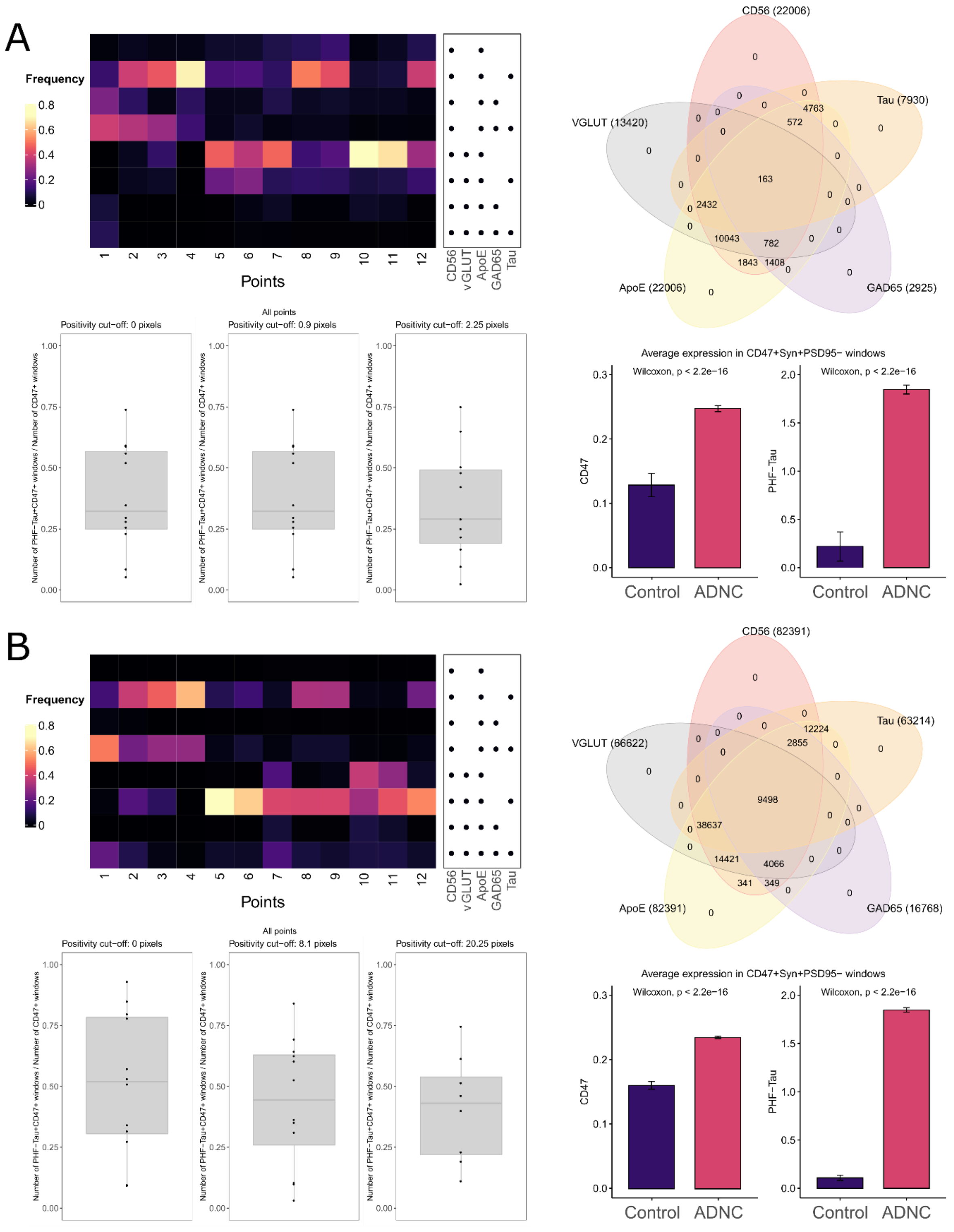


**Fig. S8. Similar MIBI analyses to Fig. 5 but with different sliding window sizes resulted in the same conclusions.** (**A**) Analyses using a window size of 3×3 (0.9 μm$^2$) and (**B**) 9×9 pixels (2.7 μm$^2$).

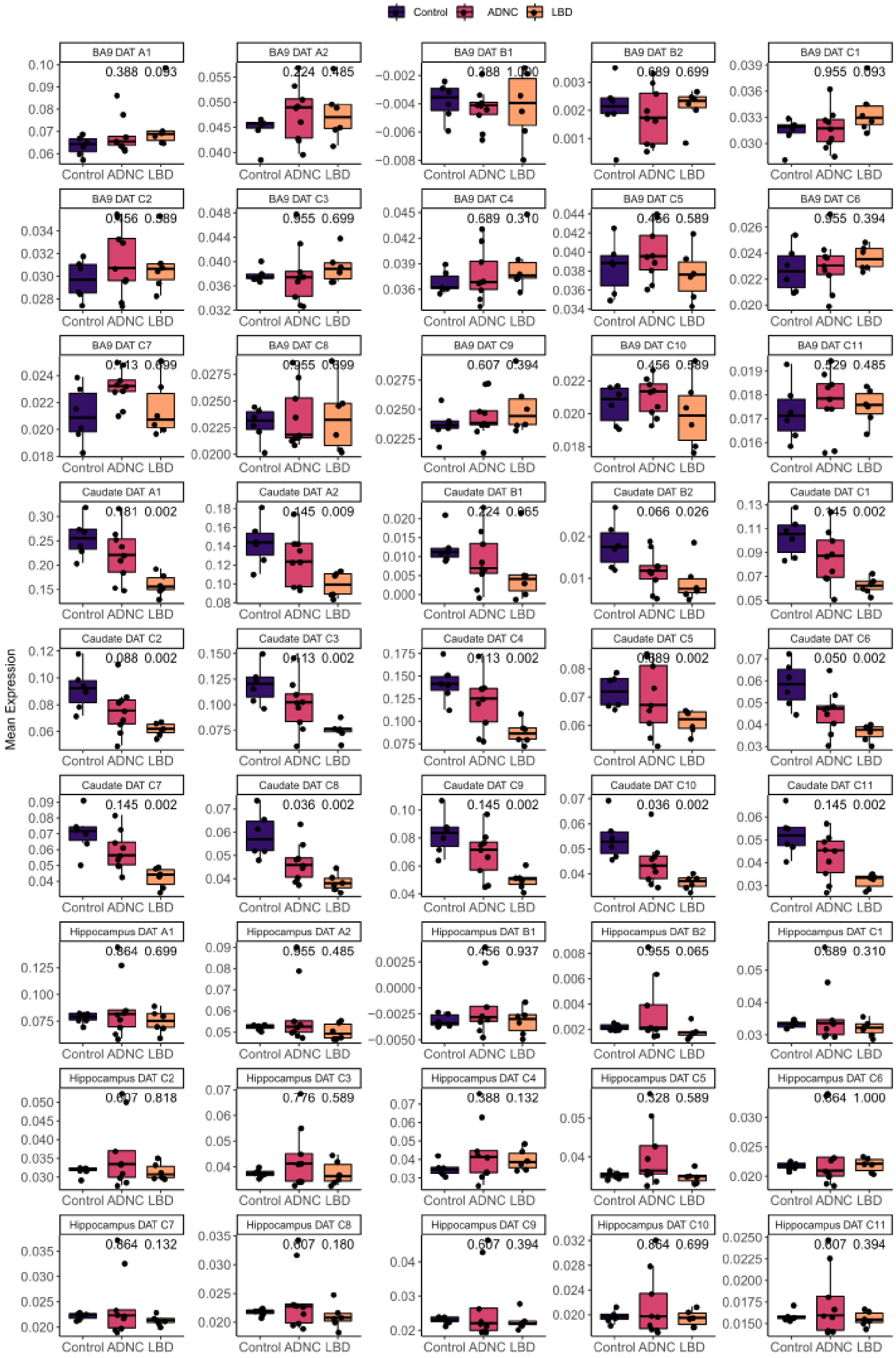

**Fig. S9**. **DAT expression in all subpopulations of the three regions conforms with traditional knowledge.** This shows that it was several folds higher in the caudate nucleus, aligning well with strong expectations from the literature.

**Supplementary Tables**

**Table S1**. Characteristics of Human Samples.

<table>
<tr><th colspan="3">Group</th><th>Control</th><th>ADNC</th><th>LBD</th></tr>
<tr><td colspan="3">N</td><td>6</td><td>9</td><td>6</td></tr>
<tr><td colspan="3">Age (yr, mean ± SD)</td><td>86 ± 8</td><td>88 ± 7</td><td>88 ± 6</td></tr>
<tr><td colspan="3">Sex (F:M)</td><td>1:2</td><td>2:1</td><td>1:1</td></tr>
<tr><td rowspan="3">Clinical Diagnosis</td><td colspan="2">Cognitively Normal (N)</td><td>6</td><td>2</td><td>2</td></tr>
<tr><td colspan="2">MCI (N)</td><td>0</td><td>1</td><td>1</td></tr>
<tr><td colspan="2">Dementia (N)</td><td>0</td><td>6</td><td>3</td></tr>
<tr><td colspan="3">PMI (hr, mean ± SD)</td><td>4.9 ± 1.6</td><td>5.0 ± 1.9</td><td>5.0 ± 0.8</td></tr>
<tr><td colspan="3">Brain (gm, mean ± SD)</td><td>1182 ± 84</td><td>1084 ± 72</td><td>1260 ± 145</td></tr>
<tr><td rowspan="9">Pathologic features^</td><td rowspan="4">ADNC (N)</td><td>No</td><td>2</td><td>0</td><td>3</td></tr>
<tr><td>Low</td><td>4</td><td>0</td><td>3</td></tr>
<tr><td>Intermediate</td><td>0</td><td>0</td><td>0</td></tr>
<tr><td>High</td><td>0</td><td>9</td><td>0</td></tr>
<tr><td rowspan="4">LB (N)</td><td>None</td><td>6</td><td>9</td><td>0</td></tr>
<tr><td>Brainstem</td><td>0</td><td>0</td><td>0</td></tr>
<tr><td>Limbic</td><td>0</td><td>0</td><td>1</td></tr>
<tr><td>Neocortical</td><td>0</td><td>0</td><td>5</td></tr>
<tr><td colspan="2">VBI (N)</td><td>None</td><td>None</td><td>None</td></tr>
</table>

| | | | | |
|---|---|---|---|---|
| | **HS (N)** | None | None | None |
| | **TDP-43(N)** | None | None | None |
| | **ARTAG (N)** | None | None | None |
| | **CTE (N)** | None | None | None |

^ Research participants who consented to brain autopsy with post mortem interval (PMI) < 8 hr had brain regions immediately processed and cryopreserved for synaptosomes as previously described *(13, 14, 19)*. Regions included Brodmann area (BA) 9 of the prefrontal cortex, hippocampus at the level of the lateral geniculate nucleus, and dorsolateral caudate nucleus. Over 5 years we collected samples from 113 brain autopsies, and included all who met the stringent clinical and pathological criteria detailed below. Individuals in the **Control** group were cognitively normal at their last clinical evaluation, which was within 2 years of death, and had cognitive test results that were in the upper three quartiles for the cohort to minimize the likelihood of interval conversion. Neuropathologic evaluation of Control individuals (n=6) had no or low AD neuropathologic change (ADNC) and no comorbidities. Individuals in the **ADNC** group (n=9) all had neuropathologic evaluations that showed only high ADNC and no evidence of any comorbidity; they were clinically diagnosed as AD dementia (n=7) or cognitively normal (n=2) or as mild cognitive impairment (n=1) within 2 years of death. Individuals in the Lewy body (LB) disease (**LBD**) group (n=6) had only limbic or neocortical LB and were clinically diagnosed as cognitively normal within 2 years of death (n=2), Parkinson's disease (PD) with MCI or dementia (n=2), or Dementia with Lewy bodies (DLB) (n=2).

**Table S2**. **SynTOF antibody panel.** Our SynTOF panel contained 38 conjugated antibodies used to probe cell type, synapse type, patho-physiological protein in synapse, protein products of several AD or PD risk genes, and markers of injury/response to injury. For quality control and normalization, we used EQ™ Four Element Calibration Beads (Fluidigm #201078), which have a unique 140Ce tag, and Cell ID™ Intercalator (191 Ir for DNA1 and 193 Ir for DNA2).

| | Antibody | Descriptor | Product identifier | Clone | Metal tag | Synaptic/ Phenotypic Marker |
|---|---|---|---|---|---|---|
| Gating | CD11b* | microglia | Fluidigm # 3148003B | M1/70 | 148 Nd | N/N |
| | MBP* | oligodendroglia | Biolegend # 808402 | SMI 99 | 150 Nd | N/N |
| | CD56 | pan-neuron | Fluidigm # 3163007B (Hu) Novus # MAB7820 (Mu) | NCAM16.2 (Hu) 809220 (Mu) | 163 Dy | Y/Y |
| | SNAP25 | pre-synaptic | Biolegend # 836304 | SMI 81 | 155 Gd | Y/Y |
| | PSD95* | post-synaptic | Biolegend # 810401 | K28/43 | 157 Gd | N/N |
| | Gephyrin* | post-synaptic | Synaptic Sys # 147011 | mAb7 | 145 Nd | N/N |
| | CD47 | pre-synaptic | BioXcell/inVivomab # BE0283 | MIAP410 | 151 Eu | Y/Y |
| | PrP | pan-neuronal | Biolegend # 800302 | 3F4 | 166 Er | Y/Y |
| | VGLUT | excitatory | Biolegend# 821301/ SySy # 135411 | N28-9/321A8 | 153 Eu | Y/Y |
| | AS | pre-synaptic | Biolegend # 807702 | LB509 | 142 Nd | Y/Y |

| | | | | | | |
|---|---|---|---|---|---|---|
| Neuron type | Tau | neuron | Millipore Sigma # MABN162 (Aka MAB361, unpurified) | TAU-5 | 172 Yb | Y/Y |
| | APP | neuron | Invitrogen # 14-9749-82 | 22C11 | 173 Yb | Y/Y |
| | GAD65 | inhibitory | Biolegend # 844502 | N-GAD65 | 169 Tm | Y/Y |
| | Calreticulin | pre-synaptic/phagocytosis | Enzo # ADI-SPA-601-F | FMC 75 | 162 Dy | Y/Y |
| | Synaptobrevin2 | pre-synaptic | Synaptic Sys # 104211 | 69.1 | 115 In | Y/Y |
| | VMAT2 | monoaminergic | Abcam # ab191121 | poly | 113 In | Y/Y |
| | SERT | serotonergic | Millipore Sigma # SAB2500950 | poly | 175 Lu | Y/Y |
| | DAT | dopaminergic | Abcam # ab5990 | hDAT-NT | 154 Sm | Y/Y |
| AD related | BIN1 | AD risk gene | Biolegend # 655602 | 99D | 147 Sm | Y/Y |
| | Aβ40 | Amyloid β peptide | Biolegend # 805402 | 11A50-B10 | 168 Er | Y/N |
| | Aβ42 | Amyloid β peptide | Biolegend #851602 | BA3-9.R | 161 Dy | Y/N |
| | PHF-tau | Paired helical filament tau | Thermo #MN1020 | AT8 | 158 Gd | Y/N |
| | ApoE | AD risk gene | Biolegend # 803303 | D6E10 | 167 Er | Y/Y |
| | TMEM230 | PD risk gene | Santa Cruz # sc-398561 | C20orf30 (G-2) | 149 Sm | Y/Y |

| | | | | | | |
|---|---|---|---|---|---|---|
| PD or DLB related | DJ1 | PD risk gene | Biolegend # 851502 | A16125E | 160 Gd | Y/Y |
| | LRRK2 | PD risk gene | Abcam # ab170993 | UDD3 30(12) | 141 Pr | Y/Y |
| | GBA1 | PD risk gene | Genzyme | 8E4 | 164 Dy | Y/Y |
| | Parkin | PD risk gene | Biolegend # 808503 | Prk 8 | 144 Nd | Y/Y |
| | p129-AS | Phospho-α-synuclein | Abcam # ab184674 | P-syn/81A | 159 Tb | Y/N |
| Injury & Response | Ac-Casp3 | apoptosis | GeneTex# GTX86909 | Poly | 146 Nd | Y/N |
| | 3-NT | oxidative stress | Sigma Millipore #05-233 | 1A6 | 165 Ho | Y/N |
| | EAAT1 | astrocyte | Thermo # PA5-72895 | poly | 089 Y | Y/N |
| | K48 | ubiquitin | Sigma Millipore # 05-1307 | Apu2 | 174 Yb | Y/N |
| | LC3B | autophagy | Novus # NB100-2220 | Poly | 171 Yb | Y/N |
| | GFAP | astrocyte | Fluidigm # 3143022B | GA5 | 143 Nd | Y/N |
| Cell Energy | GATM | creatine synthesis | Novus # NBP2-00984 | OTI1E3 | 152 Sm | Y/Y |
| | GAMT | creatine synthesis | Biorbyt # orb247514 | poly | 170 Er | Y/Y |
| | SLC6A8 | creatinine transport | mybiosource #MBS9605963 | poly | 176 Lu | Y/Y |

* The four negative markers were only used for gating presynapses, but were not included in subsequent clustering and analyses.

**Table S3**. Features (brain region, marker expression, subpopulation) that exhibited complete separation between Control vs. ADNC or Control vs. LBD.

| | **Control vs. ADNC** | | | **Control vs. LBD** | | |
|---|---|---|---|---|---|---|
| | **BA9** | **Hippocampus** | **Caudate** | **BA9** | **Hippocampus** | **Caudate** |
| **A1** | | PHF-tau, CD47 | | GFAP | | DAT |
| **A2** | | PHF-tau, CD47 | LRRK2 | GFAP | | |
| **B1** | | PHF-tau | | | | |
| **B2** | | | | | | |
| **C1** | | PHF-tau, CD47 | | | | DAT |
| **C2** | | PHF-tau | LRRK2 | | | DAT |
| **C3** | | PHF-tau, CD47 | | | | DAT |
| **C4** | | CD47 | | | | DAT |
| **C5** | | PHF-tau | | | | DAT |
| **C6** | | PHF-tau | 3NT, AS | | | DAT |
| **C7** | | PHF-tau | | | | DAT |
| **C8** | | PHF-tau | | | | DAT |
| **C9** | | PHF-tau, CD47 | | | | DAT |
| **C10** | | | PHF-tau | | | DAT |
| **C11** | | | | | | DAT |
